# GPC6 Lipid Shielding Sustains WNT5A Gradients to Prevent a RASopathy-like Developmental State

**DOI:** 10.64898/2026.09.20.752969

**Authors:** Yeonjoo Kim, Yuguang Zhao, Aurora Xinshu Xu, Asif Nakhuda, Ana-Miruna Androniciuc, Gennadiy Tenin, Bernard D. Keavney, E. Yvonne Jones, Ian J. McGough

## Abstract

Morphogen gradients require ligands to disperse across tissues from their sites of production, yet receptor engagement can consume ligand before it reaches distant cells. This poses a particular challenge for lipid-modified WNTs, whose hydrophobic palmitoleate favours ligand retention near their source and premature receptor engagement. Here we identify a palmitoleate-binding pocket in mammalian glypican-6 (GPC6), the gene mutated in the skeletal dysplasia omodysplasia, and show that shielding of the WNT5A lipid by GPC6 is required for gradient formation. Using a separation-of-function knock-in mouse allele, we demonstrate that targeted disruption of GPC6-mediated lipid shielding collapses WNT5A gradients during skeletal development by promoting premature source-proximal signalling that depletes ligand available for distal signalling. This source-proximal WNT hyperactivation engages a previously unrecognised WNT-driven RAC–PAK–MEK–ERK cascade, generating a RASopathy-like developmental state that contributes to skeletal dysplasia. Thus, GPC6 functions as a spatial gatekeeper of WNT5A utilisation, controlling the allocation of a finite ligand pool. More broadly, these findings reveal that pathway hyperactivation can arise from spatial misallocation of an endogenous signalling ligand rather than from conventional gain- or loss-of-function mutations in downstream signalling components.

## Introduction

Morphogen gradients coordinate tissue patterning, cell polarity and morphogenesis during embryonic development.^1–3^ Establishing these gradients requires secreted signalling molecules to disperse through extracellular space while remaining available for signalling at the appropriate time and place.^4,5^ Newly secreted ligands must therefore remain sufficiently mobile to spread through developing tissues. Premature receptor engagement can oppose this process by depleting the extracellular pool available for long-range distribution.^5^ How developing tissues spatially regulate the utilisation of signalling-competent ligand while preserving its availability for dispersal remains a fundamental question in developmental biology. Importantly, premature capture of ligand by signalling receptors has the potential not only to disrupt gradient formation but also to alter signalling within ligand-producing domains themselves, with consequences for embryonic development.

This challenge is particularly acute for the 19 WNT proteins found in mice and humans. WNTs carry a covalently attached palmitoleate that is essential for recognition by Frizzled (FZD) receptors and activation of downstream signalling, yet intrinsically limits extracellular solubility.^6–8^ The same lipid modification that enables signalling therefore creates a fundamental constraint on morphogen dispersal.^9,10^ By reducing extracellular mobility, the palmitoleate may favour retention of newly secreted WNT near its site of production, promoting source-proximal receptor engagement and ligand capture before tissue-scale dispersal can occur. How lipid-modified WNTs overcome this constraint to establish robust gradients remains a central unresolved question in WNT biology.

However, the palmitoleate is not only a liability for dispersal; it is also the principal determinant of Frizzled receptor engagement.^7^ By engaging the palmitoleate, molecules that mediate lipid shielding could therefore maintain WNT in a soluble, signalling-competent state while simultaneously limiting access of the exposed lipid to Frizzled receptors. If lipid shielding primarily promotes extracellular WNT solubility, its disruption would be expected to reduce signalling across the gradient. In contrast, if lipid shielding also restricts receptor capture, exposing the WNT lipid could promote premature source-proximal receptor activation and clearance of the finite extracellular ligand pool, increasing signalling near the source while depleting ligand available for tissue-scale dispersal and distal signalling.

Multiple mechanisms have been proposed to shield lipid-modified WNTs from the aqueous extracellular environment, including soluble carriers, lipoprotein particles, extracellular vesicles, specialised signalling filopodia (‘cytonemes’) that mediate direct cell–cell ligand transfer, and heparan sulfate proteoglycans.^11–18^ Glypicans, a class of cell-surface heparan sulfate proteoglycans, are particularly intriguing because our previous structural studies of the *Drosophila* glypican Dally-like protein (Dlp) revealed direct engagement of the WNT palmitoleate within a hydrophobic cavity.^18^ However, these studies did not establish whether lipid engagement regulates WNT distribution through increased extracellular solubility alone or also by limiting access of Frizzled receptors to signalling-competent ligand. Furthermore, whether glypican-mediated lipid shielding is conserved in vertebrates and regulates WNT distribution *in vivo* remains unknown, as no endogenous glypican allele specifically disrupting lipid engagement has been available. Because glypicans regulate WNT signalling through both heparan sulfate glycosaminoglycan (HS-GAG)-dependent mechanisms and direct interactions with the WNT lipid,^18–22^ separating these activities is essential for defining the specific contribution of lipid shielding to WNT organisation *in vivo*.

Among vertebrate glypicans, GPC6 provides a compelling system in which to address these questions. Loss-of-function mutations in GPC6 cause autosomal recessive omodysplasia, a skeletal dysplasia characterised by craniofacial abnormalities and impaired skeletal growth.^23,24^ Notably, several features of GPC6 deficiency overlap with developmental processes regulated by WNT5A^25–28^ and with those disrupted in Robinow syndrome, a developmental skeletal disorder resulting from mutations affecting core components of the WNT5A pathway, including WNT5A, ROR2 and DVL genes.^29–32^ Despite these phenotypic parallels, prevailing models have primarily attributed GPC6-associated skeletal defects to altered heparan sulfate-dependent Hedgehog signalling.^33^ However, GPC6 has also been implicated in WNT5A signalling in other developmental contexts,^20,34^ raising the possibility that direct binding of the WNT5A lipid by GPC6 contributes to the spatial organisation of WNT5A signalling during vertebrate skeletal development.

Here we identify a conserved lipid-binding pocket in mammalian GPC6 that directly recognises the palmitoleate moiety of WNT ligands and determine how this interaction regulates WNT availability *in vivo*. We show that disruption of GPC6 lipid shielding recapitulates major skeletal features of Gpc6 deficiency and causes collapse of WNT5A morphogen gradients. Rather than abolishing WNT5A signalling by reducing the pool of soluble, signalling-competent WNT, disruption of lipid shielding redirects signalling-competent WNT towards source-proximal receptor engagement, limiting tissue-scale ligand dispersal while driving local non-canonical WNT hyperactivation and pathological ERK signalling. These findings establish extracellular lipid shielding as a mechanism that extends beyond maintaining ligand solubility to controlling where signalling-competent WNT is utilised. More broadly, our findings demonstrate that spatial misallocation of a finite extracellular WNT pool can convert a distributed developmental signal into a localised RASopathy-like signalling state.

## Results

### Mammalian GPC6 harbours a conserved C-lobe WNT lipid-binding pocket

Our previous structural studies of *Drosophila* Dlp identified a C-lobe hydrophobic cavity formed by a bundle of α-helices that accommodates the palmitoleate moiety of WNTs.^18^ Access to this cavity is controlled by a gatekeeper residue positioned at its entrance. Across vertebrate GPC1–6 orthologues, as well as *Drosophila* Dlp and Dally, this position is highly conserved as a small side-chain residue, most commonly alanine or serine (Figure S1A).

This conservation is consistent with a requirement for unobstructed access to the lipid-binding cavity. In *Drosophila* Dlp, substitution of this serine residue with a bulky methionine hinders lipid access without perturbing overall protein fold stability,^18^ supporting its role in defining the entry point to the lipid-binding pocket. Whether this lipid-binding architecture is conserved in mammalian glypicans and mediates direct recognition of the WNT palmitoleate remains unknown.

To address this, we first examined an AlphaFold-predicted structure of human GPC6. The model revealed a characteristic glypican architecture comprising N-, M- and C-lobes. The N-lobe contains a cysteine-rich domain (CRD), and the C-lobe adopts the α-helical bundle that closely resembles the apo structure of *Drosophila* Dally-like protein (Dlp PDB ID: 3ODN).^35^ This structural similarity prompted us to model GPC6 in a lipid-bound conformation using the Dlp–lipid complex as a template (PDB ID: 6XTZ).^18^ The resulting Swiss-model structure showed close conservation of the overall C-lobe architecture and of the hydrophobic residues lining the lipid-binding cavity (Figure 1A and 1B), suggesting that the lipid-binding environment identified in Dlp is conserved in mammalian GPC6. Consistent with this prediction, a palmitoleoylated serine could be accommodated within the GPC6 cavity in a geometry closely resembling the Dlp lipid-binding mode (Figure 1B). Notably, the residue positioned at the entrance to the predicted GPC6 cavity is Ala116 in both human and mouse, corresponding to Ser168 in *Drosophila* Dlp (Figure 1B). The smaller alanine side chain is therefore compatible with unobstructed access of the palmitoleate to the hydrophobic cavity.

**Figure 1.**
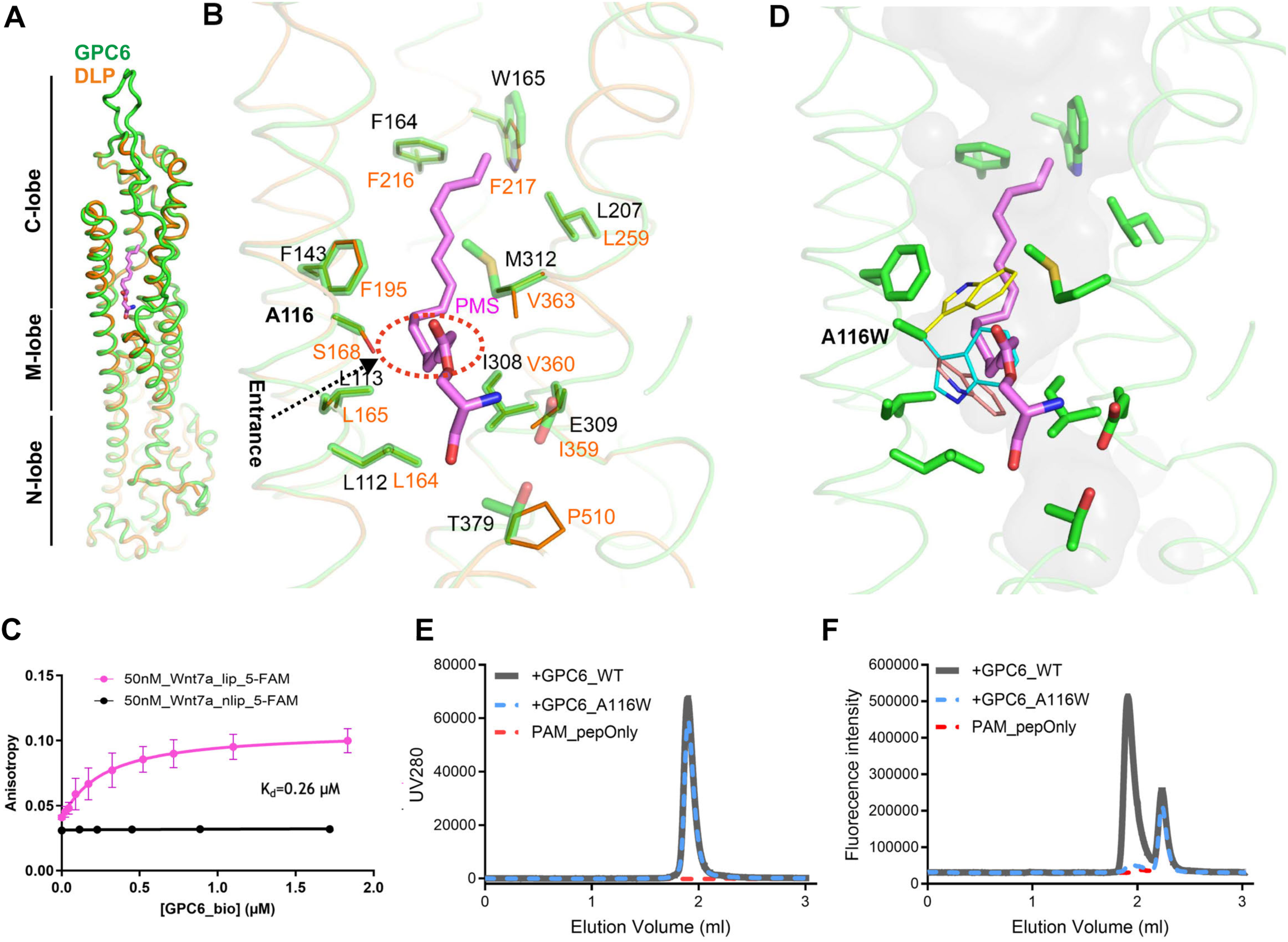
A Conserved Lipid-Binding Pocket in GPC6 Mediates WNT Lipid Recognition and Is Selectively Disrupted by a Separation-of-Function Mutation. **(A)** Cartoon representation of the AlphaFold-predicted mouse GPC6 structure (green) superimposed on the crystal structure of *Drosophila* Dally-like protein (DLP; brown; PDB 6XTZ). The palmitoleoylated serine (PMS) ligand bound to DLP is shown as purple sticks. Structural alignment reveals conservation of the putative lipid-binding pocket between vertebrate GPC6 and DLP. **(B)** Close-up view of the predicted lipid-binding pocket. Residues lining the GPC6 pocket are shown as green sticks with black labels and aligned with the corresponding DLP residues shown as brown sticks with brown labels. PMS is shown in purple. **(C)** Fluorescence anisotropy analysis of GPC6 binding to a palmitoleoylated WNT peptide. Increasing concentrations of biotinylated GPC6 ectodomain were incubated with an Alexa Fluor 488-labelled palmitoleoylated peptide derived from the WNT7A thumb region (Wnt7a_lip_5-FAM) or the corresponding non-lipidated control peptide (Wnt7a_nlip_5-FAM). Data were fitted using a one-site binding model, yielding an apparent dissociation constant (*K*_d_) of 0.26 μM for the palmitoleoylated peptide, data represent mean ± SD. **(D)** Structural model of the GPC6^A116W^ mutant. The side chain of Trp116 is shown in the three lowest-energy, non-self-clashing rotamers (yellow, cyan and salmon). Each rotamer sterically occludes the lipid-binding pocket and clashes with PMS (purple). The pocket surface is shown in transparent grey. **(E)** Fluorescence size-exclusion chromatography (FSEC) analysis of GPC6 wild-type and GPC6^A116W^. UV absorbance traces at 280 nm demonstrate comparable elution profiles for GPC6 wild-type (grey) and GPC6^A116W^ (blue dashed line), indicating that the mutation does not disrupt protein folding or overall structural integrity. The peptide-only control is shown as a red dashed line. **(F)** FSEC analysis of binding between GPC6 and a palmitoleoylated WNT peptide. Alexa Fluor 488-labelled palmitoleoylated peptide derived from the WNT7A thumb region was analysed alone (red dashed line) or following incubation with GPC6 wild-type (grey) or GPC6^A116W^ (blue dashed line). Wild-type GPC6 induces a shift in peptide elution consistent with complex formation, whereas the A116W mutation markedly reduces peptide binding.

We next tested whether mammalian GPC6 directly recognises the lipid modification of WNT ligands. Using a validated palmitoleoylated WNT7A-derived peptide corresponding to the lipidated WNT “thumb” region,^18^ fluorescence polarization assays demonstrated specific GPC6 binding, yielding an apparent equilibrium dissociation constant (*K*_d_) of 0.26 μM (Figure 1C). By contrast, the corresponding non-lipidated peptide exhibited negligible binding, demonstrating that GPC6 binding is dependent on the WNT lipid modification.

To identify a substitution that could selectively disrupt lipid binding, we focused on the conserved gatekeeper residue Ala116 at the entrance to the predicted lipid-binding cavity. Structural modelling indicated that substitution of Ala116 with the bulky residue tryptophan would occlude the cavity entrance, with Trp116 sidechain rotamers predicted to sterically clash with the palmitoleate moiety and restrict lipid access (Figure 1D). Before testing whether A116W disrupts lipid binding, we first established that the substitution did not grossly perturb GPC6 biogenesis, processing, secretion, or stability. Soluble GPC6 variants lacking the glycosylphosphatidylinositol (GPI) anchor (ΔGPI) were expressed in S2 cells and analysed in conditioned medium and cell lysates (Figure S1B). GPC6^A116W^ was secreted at levels comparable to wild-type GPC6, with no evidence of intracellular retention. In cell lysates, both variants were detected predominantly as a discrete unprocessed biosynthetic form, whereas conditioned medium contained a higher-molecular-weight, heterogeneous species consistent with GAG-modified GPC6 together with a distinct lower-molecular-weight species corresponding to a potential furin-cleaved form—a well-established consequence of glypican processing.^36,37^ The abundance and distribution of these species were comparable between wildtype GPC6 and GPC6^A116W^, indicating that the A116W substitution does not grossly impair GPC6 secretion, furin processing, or GAG modification (Figure S1B). Mutant GPC6, produced in HEK293F cells, also exhibited an identical elution profile to wild-type protein by size-exclusion chromatography, indicating no gross defect in folding or stability (Figure 1E).

Consistent with the fluorescence polarization measurements, size-exclusion chromatography demonstrated co-elution of fluorescently labelled lipidated peptide with wild-type GPC6, providing independent evidence for formation of a GPC6-WNT lipid-peptide complex (Figure 1F). In contrast, GPC6^A116W^ failed to associate with the lipid-modified WNT7A peptide despite retaining normal secretion, maturation and chromatographic behaviour (Figure 1F). Together, these data establish that the conserved GPC6 C-lobe cavity directly mediates recognition of the WNT lipid modification and identify A116W as a separation-of-function substitution that selectively abolishes lipid binding without grossly perturbing GPC6 biogenesis or stability.

### A lipid-binding-deficient *Gpc6* allele phenocopies *Gpc6* loss during skeletal development

Having established that A116W selectively disrupts lipid-dependent Wnt binding without grossly impairing GPC6 biogenesis, we next asked whether this lipid-binding activity is required for skeletal development *in vivo*. We therefore generated a *Gpc6*^A116W^ knock-in mouse carrying the endogenous A116W substitution, designed to disrupt the GPC6 lipid-binding interface. The *Gpc6*^A116W^ allele was introduced by Cas12a-mediated targeting of exon 3 followed by AAV donor-mediated homology-directed repair (HDR) (Figure S2A). This targeting strategy also generated an independent 10-bp frameshift allele (*Gpc6*^-10bp^) in exon 3. Both alleles were confirmed by sequencing (Figure S2B). Throughout, *Gpc6*^A116W^ and *Gpc6*^-10bp^ refer to homozygous *Gpc6*^A116W/A116W^ and *Gpc6*^-10bp/-10bp^ animals, respectively.

To determine whether A116W altered GPC6 expression or heparan sulfate modification in developing limbs, we analysed protein extracts from E13.5 embryonic limbs by immunoblotting. GPC6^A116W^ was expressed at levels comparable to wild-type GPC6 (Figure S2C). GPC6 protein was undetectable in *Gpc6*^-10bp^ extracts, confirming antibody specificity and supporting this allele as a functional null. The fully glycanated GPC6 species was not readily resolved under these conditions; however, heparinase treatment increased the abundance of the lower-molecular-weight GPC6 core band in both wild-type and GPC6^A116W^ samples, consistent with removal of heparan sulfate glycosaminoglycan chains (Figure S2C). qPCR analysis of E13.5 limb extracts further showed comparable *Gpc6* transcript levels between wild-type and *Gpc6*^A116W^ embryos, whereas *Gpc6* transcripts were absent in the *Gpc6*^-10bp^ frameshift allele, potentially reflecting nonsense-mediated decay (Figure S2D). Together, these data indicate that the A116W substitution does not alter GPC6 abundance or produce a detectable change in heparan sulfate modification *in vivo*. Consistent with preserved expression, GPC6 immunostaining in developing limb tissue showed a similar expression pattern in wild-type and *Gpc6*^A116W^ embryos, with strongest expression within the growth plate observed in hypertrophic and prehypertrophic zones (Figure S2E). This expression domain overlaps with the principal site of WNT5A production during skeletal development,^26^ positioning GPC6 within the ligand-producing compartment where it may regulate the extracellular organisation of WNT5A.

*Gpc6*^A116W^ mice were viable at birth (P1), in contrast to the neonatal lethality observed in the previously reported *Gpc6*^tm2b^ null mice^37^ and the *Gpc6*^-10bp^ null mice. Nevertheless, *Gpc6*^A116W^ mice were consistently smaller than wild-type littermates, as shown by whole-mount skeletal preparations, reduced body weight, and shortened tibia and fibula length (Figure 2A, 2B, 2C and 2D) recapitulating the skeletal growth defects observed in the previously reported *Gpc6^tm1Lex^* null allele and in patients with GPC6-associated omodysplasia.^23,33^ To directly compare the effects of the lipid-binding mutation with complete loss of *Gpc6*, we analysed embryos at E15.5. At this stage, *Gpc6*^A116W^ and *Gpc6*^-10bp^ embryos exhibited reduced femur and tibia length (Figure 2E and 2F), demonstrating that selective disruption of the GPC6 lipid-binding interface recapitulates the skeletal growth defects caused by GPC6 deficiency, albeit with somewhat reduced severity.

**Figure 2.**
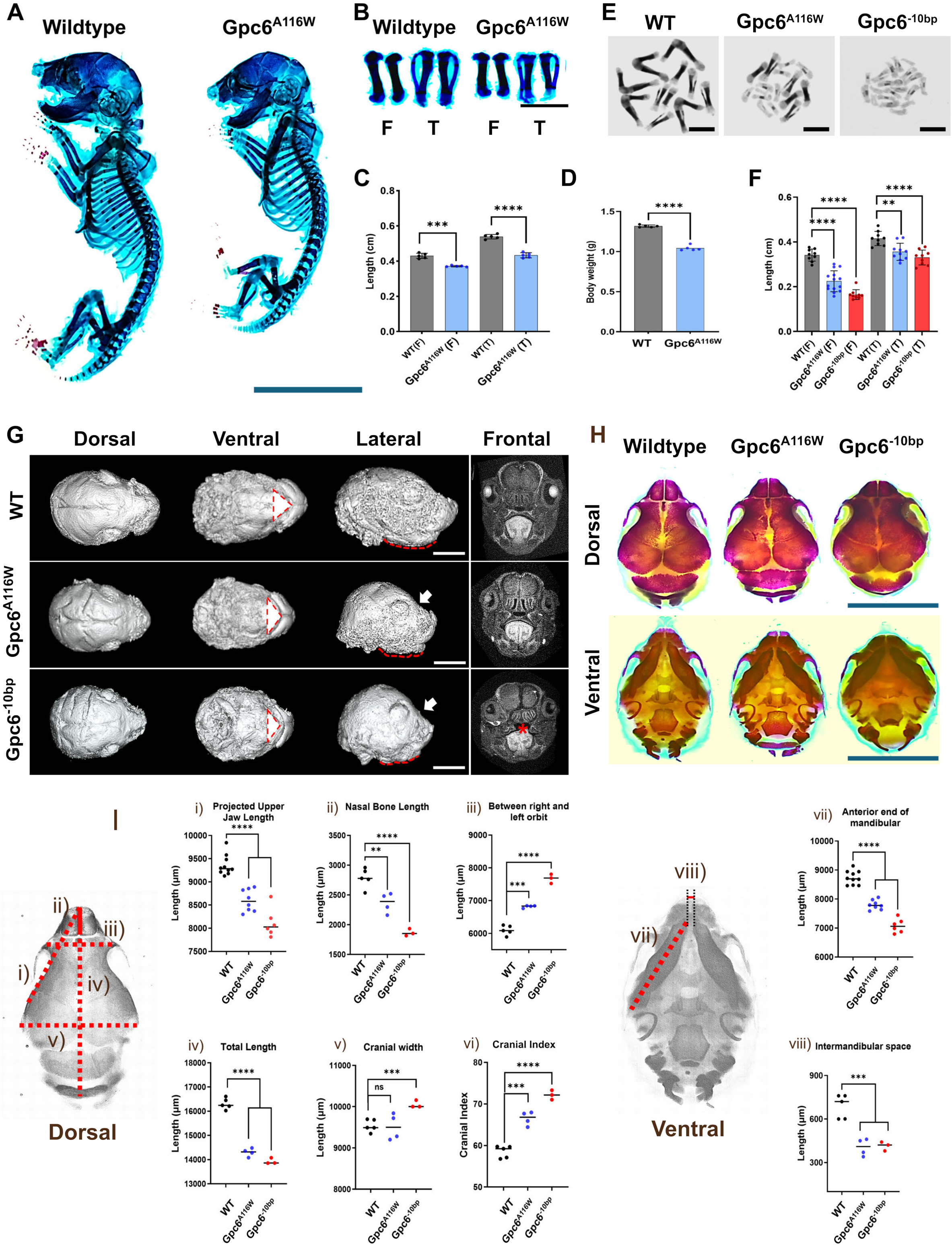
Selective Disruption of GPC6 Lipid Binding Recapitulates Gpc6 Loss- of-Function Phenotypes During Skeletal and Craniofacial Development. **(A)** Representative Alcian Blue/Alizarin Red skeletal preparations of postnatal day 1 (P1) wild-type (WT) and *Gpc6^A116W^* mice demonstrating reduced skeletal growth in mutant animals. Scale bar, 1 cm. **(B)** Representative isolated P1 femora (F) and tibiae (T) from WT and *Gpc6^A116W^*mice showing reduced long-bone growth following disruption of the GPC6 lipid-binding pocket. Scale bar, 5 mm. **(C)** Quantification of femur (F) and tibia (T) lengths from P1 WT and *Gpc6*^A116W^ mice (*n* = 5 per group; data represent mean ± SD; unpaired *t*-test with Welch’s correction, \*\*\**P* = 0.0003, \*\*\*\**P* < 0.0001). Each point represents an individual bone. **(D)** Body weight measurements of WT and *Gpc6*^A116W^ mice at P1 (*n* = 5 per group; data represent mean ± SD; unpaired *t*-test with Welch’s correction, \*\*\*\**P* < 0.0001). Each point represents an individual animal. **(E)** Representative collections of isolated femora and tibiae from E15.5 WT, *Gpc6^A116W^*, and *Gpc6^-10bp^* embryos. Scale bar, 5 mm. **(F)** Quantification of femur (F) and tibia (T) lengths from E15.5 WT, *Gpc6*^A116W^, and *Gpc6*^-10bp^ embryos [sample sizes for F: WT (*n* = 9), *Gpc6*^A116W^ (*n* = 14), *Gpc6*^-10bp^ (*n* = 9); sample sizes for T: WT (*n* = 10), *Gpc6*^A116W^ (*n* = 10), *Gpc6*^-10bp^ (*n* = 9); data represent mean ± SD; unpaired *t*-test with Welch’s correction, \*\**P* = 0.0019, \*\*\*\**P* < 0.0001]. **(G)** Three-dimensional craniofacial micro-CT reconstructions of P1 WT, *Gpc6^A116W^* and *Gpc6^-10bp^*mice shown in dorsal, ventral and lateral orientations together with representative frontal sections. Arrows indicate shortening of the snout/facial skeleton in Gpc6 mutant animals. Red asterisks indicate cleft palate, and dashed red lines indicate mandibular length. Scale bar, 200 µm. **(H)** Representative Alizarin Red-stained P1 skull preparations from WT, *Gpc6^A116W^* and *Gpc6^-10bp^* mice shown in dorsal and ventral views. Scale bar, 1 cm. **(I)** Craniofacial morphometric analyses corresponding to the measurements illustrated in the accompanying schematics: (i) projected upper jaw length (\*\*\*\**P* < 0.0001), (ii) nasal bone length (\*\*\**P* = 0.0079, \*\*\*\**P* < 0.0001), (iii) interorbital distance (\*\*\**P* = 0.0002, \*\*\*\**P* < 0.0001), (iv) total skull length (\*\*\*\**P* < 0.0001), (v) cranial width (\*\*\**P* = 0.0010), (vi) cranial index (\*\*\**P* = 0.0002, \*\*\*\**P* < 0.0001), (vii) anterior mandibular length (\*\*\*\**P* < 0.0001), and (viii) intermandibular distance (\*\*\**P* = 0.0005 (left), 0.0007 (right)). Number of mice: WT (*n* = 5), *Gpc6^A116W^* (*n* = 4), and *Gpc6^-10bp^* (*n* = 3); data represent mean ± SD; analysed by unpaired t-test with Welch’s correction.

Craniofacial analysis similarly revealed reduced anterior facial projection and mandibular hypoplasia, reflected by decreased snout length, mandible length, and overall cranial dimensions (Figure 2G, 2H and 2I). These abnormalities were evident in iodine-enhanced microCT reconstructions (Figure 2G), Alizarin Red-stained skull preparations (Figure 2H), and quantitative morphometric analyses (Figure 2I). The *Gpc6*^-10bp^ null allele exhibited pronounced craniofacial abnormalities, whereas Gpc6^A116W^ mice exhibited comparable craniofacial abnormalities in these measures, although consistently milder than those of the null allele. These phenotypes are consistent with the midfacial hypoplasia and mandibular shortening characteristic of GPC6-associated omodysplasia and WNT5A pathway-associated Robinow syndrome.^23,30,32,38^

Notably, the *Gpc6*^-10bp^ null allele recapitulated the cleft palate phenotype previously reported for the *Gpc6*^tm1Lex^ null allele,^33^ whereas cleft palate was not observed in *Gpc6*^A116W^ mice (Figure 2G). Thus, although A116W reproduces the major skeletal manifestations of GPC6 deficiency, it does not fully phenocopy complete *Gpc6* loss, indicating that some developmental functions of GPC6 are less dependent on lipid binding than skeletal growth plate development. Together, these findings demonstrate that disruption of the GPC6 lipid-binding interface is sufficient to recapitulate major skeletal manifestations of GPC6 loss *in vivo*.

### GPC6 lipid binding is required for chondrocyte proliferation, polarity and differentiation

The growth plate in developing bone is organised into sequential zones of chondrocyte differentiation that coordinate longitudinal bone growth. Resting chondrocytes give rise to proliferating chondrocytes, which undergo oriented division and form columns along the longitudinal axis of the bone, before transitioning through prehypertrophic and then hypertrophic states, the latter associated with matrix production and mineralisation.^39^ SOX9 marks the chondrogenic programme in resting, proliferating and prehypertrophic chondrocytes, but is downregulated as chondrocytes undergo hypertrophic differentiation, which is marked by expression of collagen X (COL10A1) and followed by matrix mineralisation detectable by Alizarin Red staining.^39,40^ Previous analysis of a *Gpc6* null allele identified defects in growth plate architecture, including disruption of columnar chondrocyte organisation and reduced proliferation.^33^ To directly compare the consequences of disrupting the GPC6 lipid-binding interface with complete loss of *Gpc6*, we re-examined growth plate development in *Gpc6*^tm2b^ mice, a constitutive *Gpc6* null allele,^41^ alongside *Gpc6*^A116W^ embryos. Throughout, *Gpc6*^tm2b^ mice refer to homozygous *Gpc6*^tm2b/tm2b^ animals.

Analysis of E15.5 growth plates revealed marked disorganisation of the proliferative zone in *Gpc6*^A116W^ embryos that closely resembled that observed in *Gpc6*^tm2b^ mutants, with both strains exhibiting disruption of the characteristic columnar organisation of proliferating chondrocytes observed in wild-type controls (Figure 3A). Quantification of chondrocyte morphology showed a significant reduction in cell aspect ratio in both mutant strains, indicating reduced chondrocyte elongation and flattening (Figure 3A and 3B). To quantify chondrocyte orientation, we measured the orientation of the major cellular axis relative to the longitudinal growth axis. Whereas proliferating chondrocytes in wild-type growth plates displayed a strongly aligned orientation perpendicular to the longitudinal growth axis, chondrocytes in both *Gpc6*^A116W^ and *Gpc6*^tm2b^ growth plates exhibited a broader angular distribution and reduced directional alignment (Figure 3C). We next examined the organisation of DVL2 immunofluorescence, which has previously been used as a readout of chondrocyte polarity.^42^ Rosette plots of DVL2 signal showed a pronounced directional distribution in wild-type *Gpc6* growth plates, with signal concentrated towards the two ends of the elongated chondrocyte axis, corresponding to orientations approximately perpendicular to the longitudinal growth-plate axis. This directional distribution was broader in both *Gpc6*^A116W^ and *Gpc6*^tm2b^ growth plates (Figure 3D and 3E). A similar loss of directional DVL2/3 organisation has previously been reported in *Prickle1* mutant growth plates and used as a readout of the associated loss of chondrocyte polarity.^42^ However, because DVL localisation is spatially related to the cellular axis, and the mutant chondrocytes also exhibit a marked reduction in cell elongation, the broader DVL2 distribution may in part reflect altered cell morphology. We therefore interpret the DVL2 distribution as consistent with, rather than independent evidence for, disrupted chondrocyte polarity. Equivalent disruption of growth-plate architecture was also observed in *Gpc6*^A116W^ mice at P1 (Figure S2F), demonstrating that these developmental defects persist through late embryonic and perinatal skeletal growth. These findings are reminiscent of disrupted WNT/PCP signalling in developing cartilage, as WNT5A–ROR2–VANGL2 signalling regulates chondrocyte elongation, orientation and columnar organisation during skeletal development.^25,26,42,43^

**Figure 3.**
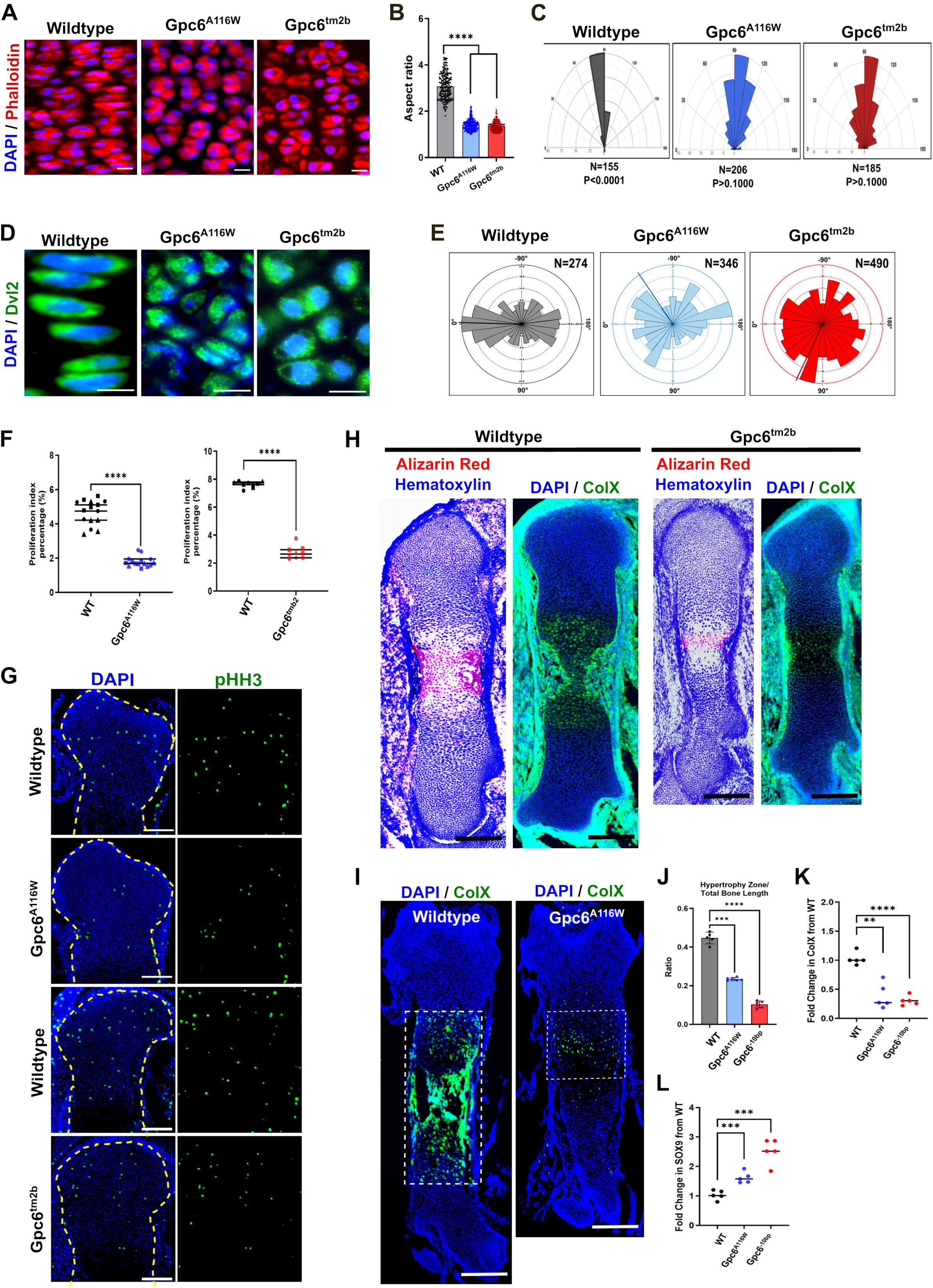
The GPC6 Lipid-Binding Interface Is Required for Growth Plate Organisation, Chondrocyte Proliferation, and Hypertrophic Differentiation. **(A)** Phalloidin staining of columnar chondrocytes within E15.5 femur growth plates from WT, *Gpc6^A116W^*and *Gpc6^tm2b^* mice revealing loss of the elongated columnar morphology characteristic of WT growth plates following disruption of GPC6 lipid binding. Scale bar 100 µm. **(B)** Quantification of chondrocyte aspect ratio within the columnar zone of E15.5 femur growth plates from WT, *Gpc6*^A116W^, and *Gpc6*^tm2b^ mice [number of analysed cells: WT (*n* = 199), *Gpc6*^A116W^ (*n* = 211), and *Gpc6*^tm2b^ (*n* = 209); data represent mean ± SD; analysed by unpaired *t*-test with Welch’s correction, \*\*\*\**P* < 0.0001]. **(C)** Semi-circular polar histograms of columnar chondrocyte orientation relative to the longitudinal growth plate axis in WT, *Gpc6^A116W^*, and *Gpc6^tm2b^* mice. Angular orientation distributions were generated using RStudio (version 2026.07.1+147) [number of analysed cells: *n* = 155 per genotype]. **(D)** Representative immunofluorescence images of DVL2 localisation in columnar chondrocytes within E15.5 femur growth plates from WT, *Gpc6^A116W^* and *Gpc6^tm2b^* mice. Scale bar 100 µm. **(E)** Circular distribution plots of DVL2 localisation demonstrating loss of polarity in *Gpc6^A116W^* and *Gpc6^tm2b^* growth plates. Plots were generated using the PCP Auto Count (PCPA) plugin in Fiji/ImageJ [analysed from 10 independent images per group; number of analysed cells: WT (*n* = 274, mean angle = −1°), *Gpc6*^A116W^ (*n* = 346, mean angle = −56°), and *Gpc6*^tm2b^ (*n* = 490, mean angle = 80°)] **(F)** Quantification of mitotic index in E15.5 femur growth plates from WT, *Gpc6^A116W^*, and *Gpc6^tm2b^* mice. Each symbol represents a biological replicate (from *n* = 3 mice per genotype). A total of 5 (left graph) and 3 (right graph) growth plate sections were analysed. Total number of analysed cells: WT (left) = 25,433, *Gpc6^A116W^* = 26,026, WT (right) = 14,173, and *Gpc6^tm2b^* = 14,011. Statistical analysis was performed using ordinary two-way ANOVA. There was a significant effect of genotype (****P < 0.0001), while the effect of experimental replicate was not significant (P = 0.9922 and 0.0605). **(G)** RepresentativePHH3 immunofluorescence staining of WT, *Gpc6^A116W^*, and *Gpc6^tm2b^* mouse growth plates used for mitotic index analysis. Scale bar 100 µm. **(H)** Representative E15.5 femur growth plate sections from WT and *Gpc6^tm2b^* embryos stained with Alizarin Red/Haematoxylin or immunolabelled for *COL10a1*. Scale bar 100 µm. **(I)** Representative *COL10a1* immunofluorescence staining of E15.5 femur growth plates from WT and *Gpc6^A116W^*embryos. Scale bar 100 µm. **(J)** Quantification of the combined hypertrophic and mineralisation zone-to-total bone length ratio in E15.5 femur growth plates from WT, *Gpc6^A116W^*, and *Gpc6^tm2b^* embryos. The ratio was calculated by dividing the combined hypertrophic and mineralising zone length by the total bone length. [number of mice: WT (*n* = 5), *Gpc6^A116W^* (*n* = 6), and *Gpc6^tm2b^*(*n* = 6); data represent mean ± SD; analysed by unpaired *t*-test with Welch’s correction, \*\*\*\**P* < 0.0001]. **(K)** Quantitative RT-PCR analysis of *Col10a1* expression. *Col10a1* transcript levels were quantified in E13.5 limb extracts from WT, *Gpc6^A116W^* and *Gpc6^tm2b^* embryos and normalised to *Gapdh*. Data are expressed as mean fold induction relative to the WT control. Error bars indicate the standard deviation (SD) of five independent experiments. Statistical significance was determined using an unpaired *t*-test with Welch’s correction (\*\**P* = 0.0011, \*\*\*\**P* < 0.0001). **(L)** Quantitative RT-PCR analysis of *Sox9* expression. *Sox9* transcript levels were quantified in E13.5 limb extracts from WT, *Gpc6^A116W^* and *Gpc6^tm2b^*embryos and normalised to *Gapdh*. Data are expressed as mean fold induction relative to the WT control. Error bars indicate the standard deviation (SD) of five independent experiments. Statistical significance was determined using an unpaired *t*-test with Welch’s correction (\*\*\**P* = 0.0008).

To determine whether these changes in growth-plate organisation were accompanied by altered chondrocyte proliferation, we quantified mitotic cells using phospho-histone H3 (pHH3) staining. Both *Gpc6*^A116W^ and *Gpc6*^tm2b^ growth plates showed a significant reduction in the mitotic index of proliferating chondrocytes relative to their respective wild-type littermate controls, with the magnitude of the reduction comparable between the two mutant strains (Figure 3F and 3G). Thus, loss of GPC6 lipid binding phenocopies the *Gpc6* null allele not only in growth-plate organisation and chondrocyte polarity-associated phenotypes, but also in the reduced proliferative output of the growth plate.

We additionally identified a previously unrecognised requirement for GPC6 in chondrocyte hypertrophic differentiation. Analysis of E15.5 femora from *Gpc6*^tm2b^ embryos revealed delayed progression of chondrocytes through hypertrophic differentiation, with COL10A1 immunostaining and Alizarin Red staining detecting only the earliest stages of hypertrophic differentiation and mineralisation, whereas substantial hypertrophic differentiation and mineralisation had already occurred in littermate controls (Figure 3H). *Gpc6*^A116W^ embryos similarly showed delayed hypertrophic differentiation, with reduced COL10A1-positive hypertrophic chondrocytes compared with wild-type littermates (Figure 3I). Quantification of the hypertrophic and early mineralising zone relative to total bone length revealed a significant reduction in both *Gpc6*^tm2b^ and *Gpc6*^A116W^ embryos, consistent with delayed progression through hypertrophic differentiation (Figure 3J). Consistent with impaired progression through the hypertrophic programme, qPCR analysis of E13.5 limb extracts revealed reduced *Col10a1* expression and persistent elevation of *Sox9* expression in both Gpc6^A116W^ and Gpc6^-10bp^ embryos (Figure 3K and 3L). As SOX9 maintains proliferative and prehypertrophic chondrocyte states and is normally downregulated during the transition to hypertrophy,^39,40^ these transcriptional changes are consistent with delayed progression of chondrocyte maturation. These data reveal a previously unrecognised requirement for GPC6 in promoting the timely transition of chondrocytes into the hypertrophic state during endochondral bone formation.

Collectively, these findings show that disruption of the GPC6 lipid-binding interface phenocopies the major growth plate defects caused by complete *Gpc6* loss, including impaired chondrocyte organisation and polarity, reduced proliferation, and delayed hypertrophic differentiation. Thus, GPC6 WNT lipid binding is a critical function underlying its role in growth plate development.

### Spatially resolved analysis reveals source-proximal non-canonical WNT hyperactivation in *Gpc6*-deficient bone

Previous studies of *Gpc6* deficiency did not identify defects in non-canonical WNT signalling in skeletal tissue and instead concluded that the skeletal phenotype arose from altered Hedgehog signalling mediated by the heparan sulfate side chains of GPC6.^33^ However, these analyses relied on whole-tissue homogenates and therefore could not assess the spatial distribution of signalling activity within the developing growth plate. As a result, localised increases and decreases in pathway activity may have been obscured in bulk measurements. Notably, the lipid-binding-deficient *Gpc6*^A116W^ allele recapitulates the major skeletal and growth-plate defects of *Gpc6* loss despite retaining intact heparan sulfate chains, indicating that disruption of HS-GAG-dependent Hedgehog regulation is unlikely to account for the major skeletal defects associated with *Gpc6* deficiency. Instead, these genetic findings implicate loss of GPC6-mediated WNT regulation as a central pathogenic mechanism. We therefore re-examined WNT pathway activity in developing long bones in GPC6 mutant alleles.

WNT5A is produced by prehypertrophic chondrocytes and regulates chondrocyte proliferation, columnar polarity and progression through the growth plate maturation programme^26^—all processes disrupted in *Gpc6*-null embryos. We therefore examined WNT5A distribution in developing long bones. Immunostaining revealed collapse of the normal WNT5A gradient in E15.5 femora from both *Gpc6*^tm2b^ and *Gpc6*^A116W^ embryos (Figure 4A and 4B), while Western blot analysis of E13.5 limb lysates confirmed reduced total WNT5A protein levels in both mutants (Figure 4C). qPCR analysis of E13.5 limb extracts revealed that *Wnt5a* transcript levels were not reduced in either mutant (Figure 4D), indicating that the reduction in WNT5A protein is not transcriptionally mediated. One possibility is that loss of GPC6-mediated lipid shielding reduces the extracellular stability of

**Figure 4.**
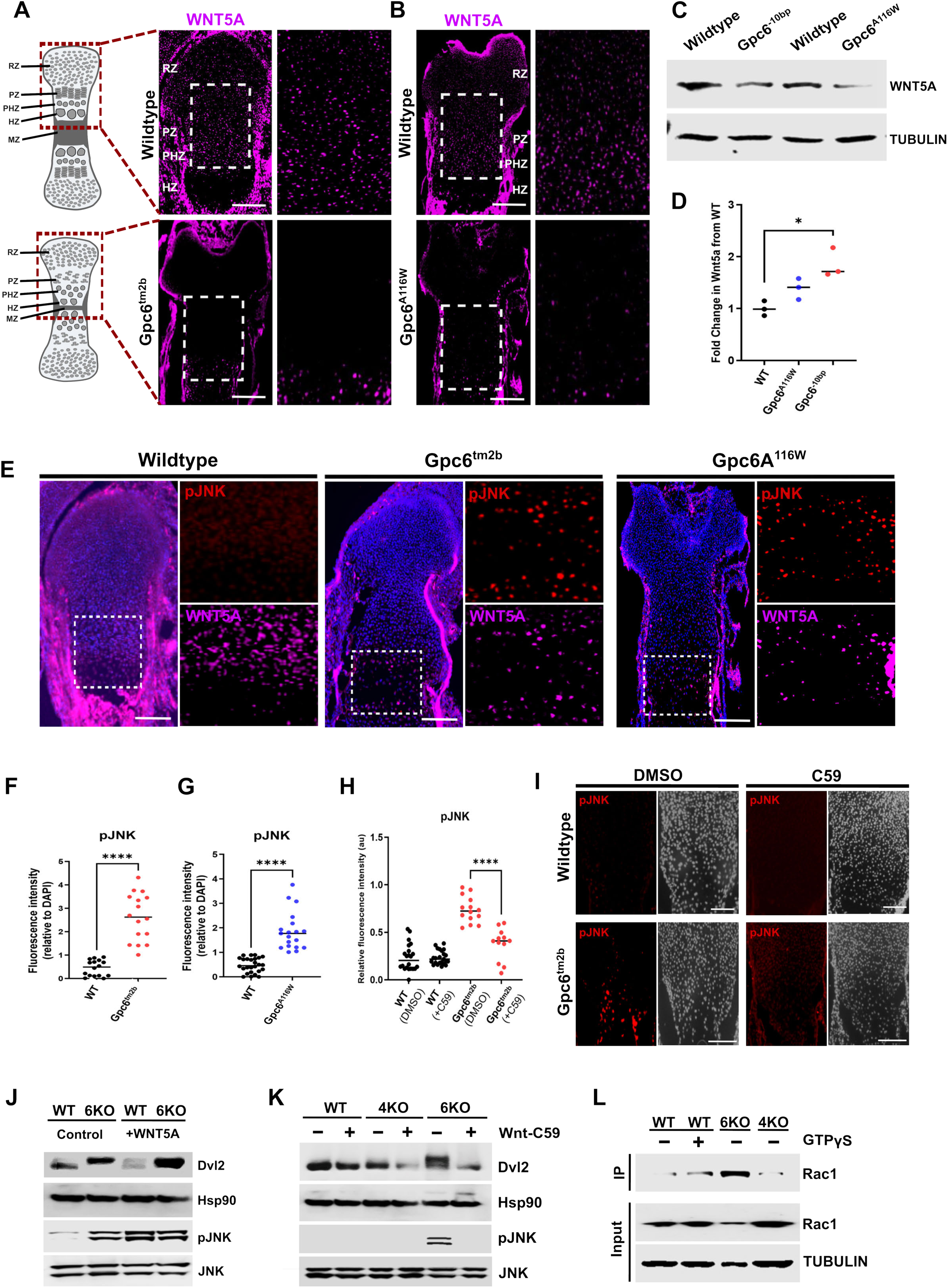
WNT5A Gradient Collapse Causes Source-Proximal WNT Hyperactivation in Gpc6-Deficient Bone. **(A)** Schematic illustrating growth plate organisation and the region analysed for spatial WNT5A distribution (left). Representative WNT5A immunofluorescence images of E15.5 femora from WT and *Gpc6^tm2b^* littermates (right) showing collapse of the physiological WNT5A gradient in *Gpc6^tm2b^*growth plates. Scale bar 100 µm. RZ, resting zone; PZ, proliferative zone; PHZ, prehypertrophic zone; HZ, hypertrophic zone; MZ, mineralization zone. **(B)** Representative WNT5A immunofluorescence images of E15.5 femora from WT and *Gpc6^A116W^* littermates showing collapse of the physiological WNT5A gradient in *Gpc6^A116W^*growth plates. Scale bar 100 µm. **(C)** Western blot analysis of WNT5A protein abundance in E13.5 limb extracts from *Gpc6^tm2b^*and *Gpc6*^A116W^ embryos and corresponding wild-type littermates, showing reduced total WNT5A protein levels in both mutants. **(D)** Quantitative RT-PCR analysis of *Wnt5a* expression. *Wnt5a* transcript levels were quantified in E13.5 limb extracts from WT, *Gpc6^A116W^* and *Gpc6^tm2b^* embryos and normalised to *Gapdh*. Data are expressed as mean fold induction relative to the WT control. Error bars indicate the standard deviation (SD) of five independent experiments. Statistical significance was determined using an unpaired *t*-test with Welch’s correction (\**P* = 0.0193). **(E)** Representative pJNK immunofluorescence images of E15.5 femoral growth plates from WT, *Gpc6^tm2b^*, and *Gpc6^A116W^* embryos. Insets show the WNT5A-producing prehypertrophic domain, revealing increased pJNK signal in both mutant genotypes. WNT5A images from mutant samples were acquired at higher detector settings than wild-type samples to visualise residual WNT5A signal within the prehypertrophic domain and permit comparison with pJNK localisation. Scale bar 100 µm. **(F)** Quantification of pJNK signal intensity in the prehypertrophic domain. Relative fluorescence intensity of pJNK was measured within the WNT5A-producing prehypertrophic domains of growth plates from WT and *Gpc6^tm2b^* (*n* = 16 sections per group) embryos. Signals were normalised to DAPI fluorescence intensity. The significant increase in pJNK signal demonstrates elevated non-canonical WNT pathway activity in *Gpc6^tm2b^* growth plates relative to wild-type controls. Statistical significance was determined using an unpaired t-test with Welch’s correction (\*\*\*\**P* < 0.00001). **(G)** Quantification of pJNK signal within the WNT5A-producing prehypertrophic domain demonstrating increased non-canonical WNT pathway activity in *Gpc6^A116W^*growth plates relative to wild-type controls. Relative fluorescence intensity of pJNK was measured within the WNT5A-producing prehypertrophic domains of growth plate sections from WT (*n* = 26 sections) and *Gpc6^A116W^*(*n* = 19 sections) embryos. Signals were normalised to DAPI fluorescence intensity. The significant increase in pJNK signal demonstrates elevated non-canonical WNT pathway activity in *Gpc6^A116W^* growth plates relative to wild-type controls. Statistical significance was determined using an unpaired t-test with Welch’s correction (\*\*\*\**P* < 0.00001). **(H)** Quantification of pJNK signal intensity following Porcupine inhibition. Femoral explants from WT and *Gpc6^tm2b^* embryos were cultured for 24 hours in the presence of either vehicle (DMSO) or the Porcupine inhibitor C59 (50 nM). Relative fluorescence intensity of pJNK was measured within the WNT5A-producing prehypertrophic domains of growth plate sections from WT + DMSO (*n* = 22), WT + C59 (*n* = 23), *Gpc6^tm2b^* + DMSO (*n* = 14), and *Gpc6^tm2b^*+ C59 (*n* = 12) sections. Signals were normalised to DAPI fluorescence intensity. Porcupine inhibitor C59 significantly suppresses the elevated pJNK activity observed in Gpc6-deficient bone. Statistical significance was determined using an unpaired t-test with Welch’s correction (\*\*\*\**P* < 0.00001). **(I)** Representative pJNK immunofluorescence images of paired E14.*5 Gpc6^tm2b^* femoral explants cultured for 24 h in DMSO or the Porcupine inhibitor C59, showing loss of elevated pJNK signal following inhibition of WNT secretion. Scale bar 100 µm. **(J)** Immunoblot analysis of DVL2 and JNK phosphorylation in wild-type and Gpc6-deficient ATDC5 cells cultured in control or WNT5A-conditioned L-cell medium for 30 min, showing elevated basal non-canonical WNT pathway activity in Gpc6-deficient cells. **(K)** Immunoblot analysis of DVL2 and JNK phosphorylation in WT, Gpc4-deficient, and Gpc6-deficient ATDC5 cells treated with DMSO or the Porcupine inhibitor C59, showing suppression of elevated pathway activity in Gpc6-deficient cells following inhibition of WNT secretion. **(L)** RAC1-GTP pull-down assay in WT, Gpc4-deficient, and Gpc6-deficient NIH-3T3 cells demonstrating increased RAC1-GTP loading selectively in Gpc6-deficient cells.

WNT5A, leading to increased ligand loss before it can disperse through the tissue. However, reduced steady-state ligand abundance could also reflect increased receptor-mediated utilisation if signalling-competent WNT5A is captured and consumed near its site of production. We therefore asked whether the reduction in WNT5A abundance was associated with diminished or increased local pathway activity.

Phosphorylated JNK (pJNK) is an established downstream readout of WNT5A-dependent non-canonical signalling.^44^ Analysis of pJNK revealed a marked increase in signalling within the WNT5A-producing prehypertrophic domain in Gpc6^tm2b^ embryos, overlapping the residual WNT5A expression, despite the global reduction in WNT5A abundance (Figure 4E and 4F). Thus, loss of GPC6 does not simply diminish WNT5A signalling but instead produces a localised state of pathway hyperactivation within the ligand-producing domain. Strikingly, a similar increase in pJNK was observed in the WNT5A-producing domain of *Gpc6*^A116W^ embryos (Figure 4E and 4G). This demonstrates that localised pathway hyperactivation is recapitulated by selective disruption of the GPC6 lipid-binding interface, linking defective WNT5A lipid engagement directly to the altered spatial organisation of signalling.

To determine whether this localised pathway activation is a tissue-intrinsic consequence of *Gpc6* loss and whether it depends on endogenous WNT secretion, we established an ex vivo bone explant assay that preserves the native spatial organisation and cellular interactions of the growth plate while removing systemic influences and permitting direct experimental manipulation of WNT secretion. Paired E14.5 femora from *Gpc6*^tm2b^ embryos were cultured for 24 h in either DMSO or the Porcupine inhibitor C59, which blocks secretion of newly synthesised WNT ligands.^45^ C59 treatment reduced the elevated pJNK signal, demonstrating that the increased pathway activity is dependent on endogenous WNT secretion and persists in the absence of systemic influences (Figure 4H and 4I).

Thus, loss of GPC6 produces a spatially paradoxical WNT phenotype: WNT5A abundance and tissue-scale distribution are reduced, yet WNT-dependent signalling is increased near the ligand source. Selective disruption of the GPC6 lipid-binding interface reproduces both features, supporting a model in which GPC6-mediated lipid shielding limits source-proximal utilisation of signalling-competent WNT and thereby preserves ligand for tissue-scale distribution.

### Loss of *Gpc6* establishes an endogenous autocrine non-canonical WNT signalling state that drives chronic RAC activation

To further dissect the mechanisms underlying pathway activation and enable analysis of downstream signalling events that are difficult to resolve *in vivo*, we established an *in vitro* model using the well-characterised ATDC5 mouse chondrogenic cell line, which can be induced to undergo hypertrophic differentiation.^46,47^ qPCR analysis confirmed expression of both *Gpc6* and the closely related glypican *Gpc4* in ATDC5 cells (Figure S3A). Given their overlapping expression patterns and reported functional similarities,^48,49^ we generated independent *Gpc6*- and *Gpc4*-knockout cell lines to define their individual contributions to non-canonical Wnt signalling (Figure S3B and S3C).

To determine how loss of GPC6 affects cellular responses to WNT5A, including the possibility of altered pathway sensitivity, we examined WNT5A-dependent signalling in ATDC5 cells. Treatment of wild-type ATDC5 cells with WNT5A-conditioned medium for 30 min induced JNK phosphorylation together with a characteristic mobility shift of DVL2,^50^ a core intracellular transducer of WNT receptor signalling, consistent with DVL2 phosphorylation and activation of non-canonical WNT signalling (Figure 4J). These responses established that ATDC5 cells are competent to activate the non-canonical WNT pathway in response to WNT5A stimulation. In contrast, *Gpc6*-deficient cells exhibited constitutively elevated phosphorylation of both DVL2 and JNK in the absence of exogenous WNT5A (Figure 4J). Treatment with the Porcupine inhibitor C59 abolished this constitutive activation, demonstrating that it depends on endogenous WNT secretion and indicating increased autocrine WNT pathway activity (Figure 4K). By comparison, *Gpc4*-deficient cells showed no detectable increase in pathway activation under the same conditions (Figure 4K). Together, these findings indicate that loss of GPC6 selectively increases WNT-dependent pathway activity, resulting in constitutive activation of non-canonical WNT signalling driven by endogenous WNT.

To ask if GPC6 deficiency also leads to increased non-canonical WNT signalling beyond chondrogenic cells, we performed complementary experiments in NIH-3T3 fibroblasts, which express GPC3, GPC4, GPC5 and GPC6 (Figure S3D). Independent *Gpc4*- and *Gpc6*-knockout NIH-3T3 cell lines were generated and validated (Figure S3E and S3F). Consistent with our findings in ATDC5 cells, loss of *Gpc6*, but not *Gpc4*, increased the phosphorylation-dependent mobility shifts of both DVL2 and the non-canonical WNT co-receptor ROR2 (Figure S3G), which undergoes phosphorylation-dependent mobility shifts in response to WNT5A stimulation,^28^ indicating enhanced activation of the non-canonical WNT pathway.

These changes occurred in the absence of exogenous WNT ligand. Concordant with increased pathway activation, *Gpc6*, but not *Gpc4*, loss also increased loading of RAC1-GTP—a downstream effector of non-canonical WNT signalling^51^ (Figure 4L)—thus placing enhanced RAC activation downstream of the autocrine signalling state. Notably, these phenotypes occurred despite the expression of multiple other glypicans, indicating that they do not detectably compensate for GPC6 in restraining endogenous non-canonical WNT signalling under these conditions.

### RAC–PAK hyperactivation couples non-canonical Wnt signalling to ERK activation

The effect of GPC6 loss on RAC1-GTP loading was particularly intriguing in light of previous evidence that RAC signalling dosage influences chondrocyte maturation.^52^ We therefore asked whether this elevated RAC1 signalling could provide a mechanistic link between dysregulated non-canonical WNT signalling and the skeletal abnormalities caused by GPC6 loss. Beyond its cytoskeletal functions, RAC can act as a potent upstream activator of PAK-dependent ERK1/2 signalling,^53–58^ and oncogenic RAC1 signalling contributes to pathological ERK1/2 activation in multiple tumour settings.^59–61^ Given the established role of sustained ERK1/2 activation in growth plate dysregulation and RASopathy-associated skeletal disease,^62–66^ we asked whether increased RAC signalling following *Gpc6* loss engages the ERK1/2 pathway.

Consistent with increased RAC1-GTP loading, *Gpc6*-deficient ATDC5 cells exhibited enhanced PAK1 phosphorylation at Ser199/Ser204, autophosphorylation sites,^67,68^ accompanied by increased phosphorylation of MEK at both the PAK-responsive regulatory site (Ser298)^53,54^ and its activation loop residues, together with elevated ERK1/2 activation loop phosphorylation (Figure 5A and 5B). ERK1/2 activation was abolished by C59 treatment, demonstrating dependence on endogenous WNT secretion (Figure 5B). Thus, loss of *Gpc6* engages a RAC–PAK–MEK–ERK signalling cascade downstream of an endogenous autocrine WNT signalling state.

**Figure 5.**
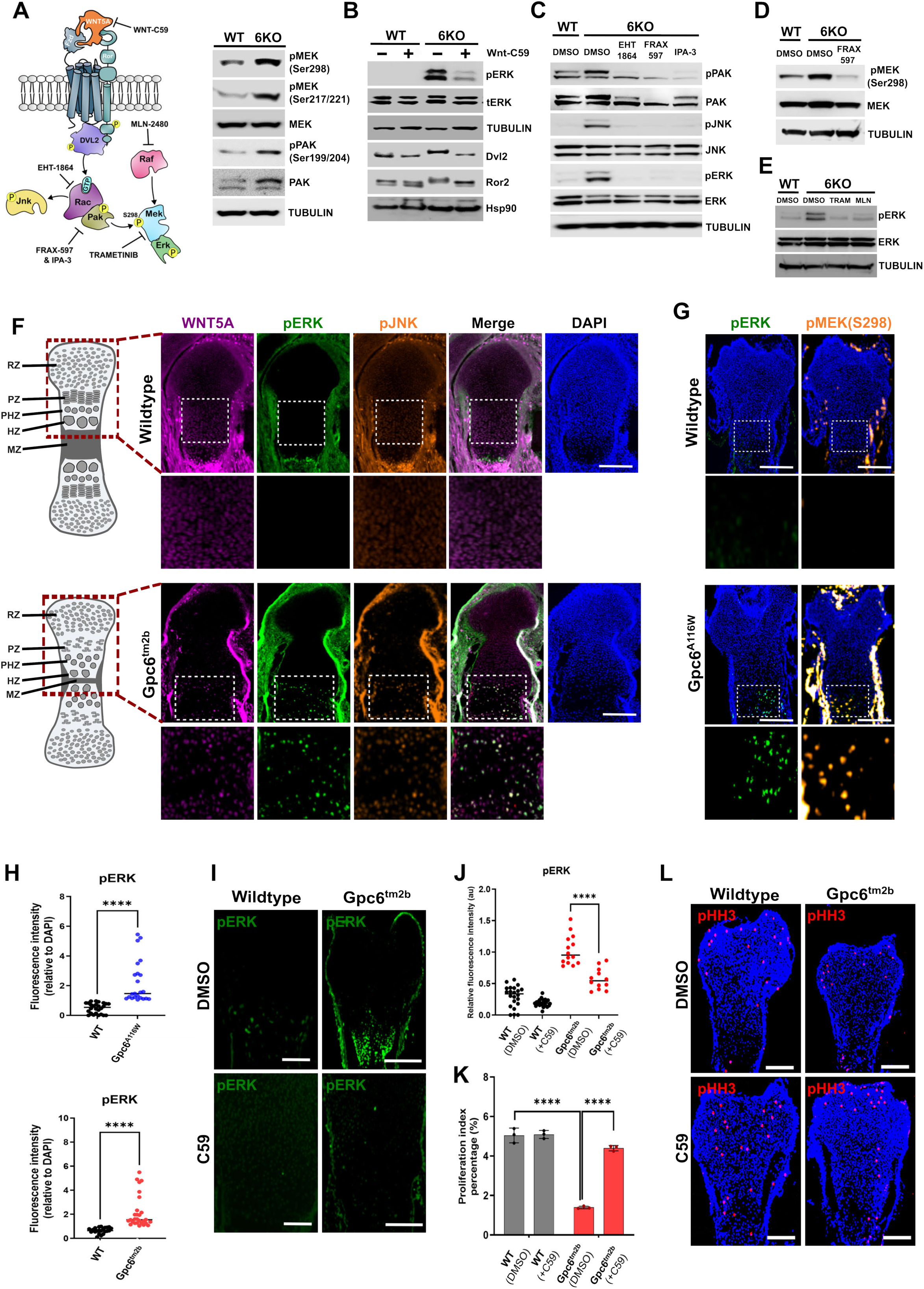
WNT-RAC-PAK Signalling Drives ERK Hyperactivation Following Gpc6 Loss. **(A)** Schematic of the proposed WNT5A-RAC-PAK-MEK1-ERK signalling pathway and sites of pharmacological inhibition (left). Immunoblot analysis of PAK1 phosphorylation at Ser199/Ser204 and MEK phosphorylation at Ser298 and Ser217/221 in WT and Gpc6-deficient ATDC5 cells (right). **(B)** Immunoblot analysis of DVL2 and ROR2 phosphorylation-dependent mobility shifts together with ERK1/2 activation-loop phosphorylation (Thr202/Tyr204 in ERK1 and Thr185/Tyr187 in ERK2) in WT and Gpc6-deficient ATDC5 cells treated with DMSO or the Porcupine inhibitor C59 for 16 h. **(C)** Immunoblot analysis of PAK1, JNK and ERK phosphorylation in Gpc6-deficient ATDC5 cells following 3 h treatment with the RAC inhibitor EHT1864 or the PAK inhibitors FRAX597 and IPA-3. **(D)** Immunoblot analysis of MEK phosphorylation at the PAK-responsive regulatory site Ser298 in Gpc6-deficient ATDC5 cells following 3 h treatment with the PAK inhibitor FRAX597. **(E)** Immunoblot analysis of ERK phosphorylation in Gpc6-deficient ATDC5 cells following 3 h treatment with the MEK inhibitor trametinib or the pan RAF inhibitor MLN2480. **(F)** Schematic illustrating growth plate organisation together with representative immunofluorescence images of WNT5A, pJNK and pERK in E15.5 femoral growth plates from WT and *Gpc6^tm2b^*embryos. Insets show the WNT5A-producing prehypertrophic domain. Scale bar 100 µm. **(G)** Representative immunofluorescence images of pERK and pMEK(S298) in E15.5 femoral growth plates from WT and *Gpc6^A116W^* embryos and corresponding wild-type littermates. Scale bar 100 µm. **(H)** Quantification of pERK signal intensity in the prehypertrophic domain. Relative fluorescence intensity of pERK was measured within the WNT5A-producing prehypertrophic domains of growth plate sections from WT and *Gpc6^A116W^*embryos (above; *n* = 26 and *n* = 23 sections, respectively) and WT and *Gpc6^tm2b^* embryos (below; *n* = 22 and *n* = 28 sections, respectively). Signals were normalised to DAPI fluorescence intensity. The significant increase in pERK signal demonstrates elevated MAPK pathway activity in *Gpc6^tm2b^* growth plates relative to WT controls. Statistical significance was determined using an unpaired t-test with Welch’s correction (****P < 0.00001). **(I)** Representative pERK immunofluorescence images of paired E14.5 *Gpc6^tm2b^*femoral explants cultured for 24 h in DMSO or the Porcupine inhibitor C59. Scale bar 100 µm. **(J)** Quantification of pERK signal intensity following Porcupine inhibition. Femoral explants from WT and *Gpc6^tm2b^* embryos were cultured for 24 hours in the presence of either vehicle (DMSO) or the Porcupine inhibitor C59 (50 nM). Relative fluorescence intensity of pERK was measured within the WNT5A-producing prehypertrophic domains of growth plate sections from WT + DMSO (*n* = 22), WT + C59 (*n* = 23), *Gpc6^tm2b^* + DMSO (*n* = 14), and *Gpc6^tm2b^*+ C59 (*n* = 12) sections. Signals were normalised to DAPI fluorescence intensity. Porcupine inhibition significantly suppresses the elevated pERK activity observed in Gpc6-deficient bone. Statistical significance was determined using an unpaired t-test with Welch’s correction (\*\*\*\**P* < 0.00001). **(K)** Quantification of mitotic index following Porcupine inhibition. Femoral explants from WT and *Gpc6^tm2b^* embryos were cultured for 24 hours in the presence of either vehicle (DMSO) or the Porcupine inhibitor C59 (50 nM) across three independent replicates (*n* = 3 explants per group). The mitotic index was determined from a total analysed population of 13,502 cells for WT + DMSO, 12,025 cells for WT + C59, 9,316 cells for *Gpc6^tm2b^* + DMSO, and 14,598 cells for *Gpc6^tm2b^* + C59. Data demonstrate that Porcupine inhibition restores the altered mitotic index observed in *Gpc6*-deficient bone. Statistical significance was determined by ordinary two-way ANOVA followed by Šídák’s multiple comparisons test to evaluate pre-selected pairs (WT *vs.* KO, and KO *vs.* KO+C59). Genotype and C59 treatment effects were highly significant (\*\*\*\**P* < 0.0001), while the replicate effect was not significant. **(L)** Representative pHH3 immunofluorescence staining of DMSO- and C59-treated E14.5 *Gpc6^tm2b^* femoral explants. Scale bar 100 µm.

To determine whether activation of the RAC–PAK–MEK–ERK cascade following *Gpc6* loss extended beyond chondrogenic cells, we analysed NIH-3T3 fibroblasts lacking either *Gpc6* or *Gpc4*. Consistent with the increased non-canonical WNT signalling observed following *Gpc6* loss in these cells, *Gpc6* deficiency resulted in increased phosphorylation of PAK1, MEK and ERK1/2, whereas loss of *Gpc4* failed to activate these pathway components (Figure S3G). These findings further demonstrate that GPC6, but not GPC4, has a non-redundant role in restraining autocrine non-canonical WNT–RAC–PAK–MEK–ERK signalling.

To define the pathway hierarchy linking aberrant WNT activation to MAPK signalling, we pharmacologically inhibited RAC and PAK. Treatment with the RAC inhibitor EHT1864^69^ reduced phosphorylation of PAK1 and JNK and suppressed ERK1/2 activation in Gpc6-deficient cells, consistent with RAC acting downstream of non-canonical WNT signalling and upstream of MAPK activation (Figure 5C). Similarly, inhibition of PAK1 using either the catalytic inhibitor FRAX597^70^ or the allosteric inhibitor IPA-3^71^ reduced PAK1 phosphorylation and suppressed ERK1/2 activation (Figure 5C). PAK inhibition also reduced phosphorylation of the PAK target site MEK Ser298, confirming that this residue is regulated by PAK activity in this context (Figure 5D). Together, these findings establish a RAC–PAK signalling module as a critical link coupling aberrant non-canonical WNT signalling to downstream MEK–ERK activation following *Gpc6* loss.

To further define the signalling pathway linking non-canonical WNT activation to ERK1/2, we inhibited MEK and RAF using trametinib^72^ and tovorafenib (MLN2480),^73^ respectively. Both inhibitors abolished the elevated ERK1/2 phosphorylation observed in *Gpc6*-deficient cells (Figure 5E). Importantly, the ability of RAF inhibition to suppress ERK1/2 activation indicates that RAC–PAK signalling does not bypass the canonical RAF–MEK–ERK cascade but rather potentiates its signalling output.

Finally, we sought to determine which endogenous WNT ligand drives pathway activation following *Gpc6* loss. WNT5A was a strong candidate given its prominent expression within the developing growth plate and the marked disruption of WNT5A abundance and gradient architecture observed in *Gpc6*-deficient long bones. However, because *Wnt5b* is also expressed by chondrocytes and has overlapping expression and functions during skeletal development,^26^ we simultaneously suppressed both ligands using pooled siRNAs. Knockdown of *Wnt5a* and *Wnt5b* reduced non-canonical WNT pathway activation in *Gpc6*-deficient cells, as evidenced by diminished ROR2 mobility shift, reduced phosphorylation of the PAK target site MEK Ser298, and suppression of ERK1/2 phosphorylation (Figure S3H). These findings establish endogenous WNT5-family ligands as the drivers of RAC–PAK– MEK–ERK activation following *Gpc6* loss.

Together, these findings identify sustained ERK1/2 activation as a previously unrecognised consequence of *Gpc6* loss. Aberrant ERK1/2 signalling is driven by endogenous WNT ligand and requires RAC–PAK–MEK signalling, establishing a mechanistic link between defective non-canonical WNT regulation and activation of the RASopathy-associated ERK1/2 pathway. We therefore examined whether ERK1/2 signalling is similarly elevated within the developing skeleton *in vivo*.

### GPC6 lipid binding restrains WNT-dependent ERK1/2 activation near the WNT source in the developing growth plate

To determine whether aberrant ERK1/2 activation accompanies WNT5A gradient collapse *in vivo*, we performed triple immunofluorescence for WNT5A, phosphorylated JNK (pJNK) and phosphorylated ERK1/2 (pERK) in developing long bones. This revealed a marked increase in ERK1/2 activation in *Gpc6*-null embryos, specifically within the WNT5A-producing prehypertrophic chondrocyte domain (Figure 5F and 5H). These same cells also exhibited elevated pJNK, indicating that enhanced non-canonical WNT and ERK1/2 signalling converge within the prehypertrophic zone *in vivo*. Consistent with the signalling mechanism defined *in vitro*, Gpc6-null growth plates likewise exhibited increased phosphorylation of MEK at the PAK-responsive Ser298 site within this region (Figure S4A), providing *in vivo* evidence for activation of the RAC–PAK–MEK–ERK signalling axis upstream of ERK1/2.

Importantly, the same spatially restricted increase in pERK and pMEK(S298) was observed in *Gpc6*^A116W^ embryos, with elevated signalling concentrated within the prehypertrophic domain (Figure 5G and 5H). These findings indicate that the WNT lipid-binding function of GPC6 is required to restrain WNT-dependent ERK1/2 activation at the WNT5A source.

This localised signalling response was not restricted to the femoral growth plate. Similar spatially restricted increases in pJNK and pERK were observed in the tibial growth plate of *Gpc6*-deficient embryos and coincided with collapse of the WNT5A gradient (Figure S4B and S4C). These findings indicate that WNT5A gradient collapse and source-proximal activation of non-canonical WNT and ERK1/2 signalling are general consequences of Gpc6 loss during endochondral skeletal development.

To establish whether this ERK-active state was driven by endogenous WNT secretion and was intrinsic to developing skeletal tissue, we performed *ex vivo* bone explant cultures. E14.5 femora were isolated and cultured in the presence of C59. The elevated pERK signal observed in *Gpc6*-deficient growth plates was maintained in isolated bone explants but reduced following inhibition of WNT secretion (Figure 5I and 5J), demonstrating that ERK activation is dependent on endogenous WNT secretion and does not require systemic inputs. Notably, C59 treatment also increased pHH3 staining in Gpc6-deficient femora (Figure 5K and 5L), suggesting that excessive local WNT signalling contributes to the reduced proliferative state associated with loss of GPC6. Together, these findings demonstrate that loss of GPC6 generates a tissue-autonomous, WNT-dependent ERK-active state within the developing skeleton.

### GPC6 expression is spatially coupled to WNT5A-producing domains across developing tissues

Having established that GPC6 regulates WNT5A distribution during skeletal development, we asked whether this relationship represents a broader developmental principle. Examination of additional embryonic tissues revealed striking spatial concordance between GPC6 and WNT5A expression domains. In developing craniofacial structures, including tooth buds, mandible, whisker follicles, tongue and pituitary, GPC6 expression closely overlapped with regions expressing WNT5A (Figure S5). Consistent with this association, previous developmental studies identified GPC6 expression within WNT5A-producing intestinal stromal populations.^34^ Together, these observations reveal a recurrent developmental relationship between GPC6 and WNT5A-producing cells across diverse organ systems. This conserved spatial coupling suggests that GPC6 may serve as a general extracellular buffering mechanism for lipid-modified WNT5A, allowing ligand dispersal away from producing cells while preventing accumulation of signalling-competent WNT within the source domain.

### Loss of GPC6 delays chondrocyte differentiation via ERK-dependent signalling

To determine whether ERK1/2 hyperactivation contributes to the delayed chondrocyte maturation associated with *Gpc6* loss, we monitored hypertrophic differentiation of ATDC5 cells over a 35-day time course under mineralising culture conditions. Sustained ERK1/2 activation is known to impair chondrocyte maturation and delay hypertrophic differentiation,^62^ including in RASopathy models driven by gain-of-function mutations in PTPN11/SHP2.^66^ We therefore asked whether a similar ERK1/2-dependent mechanism operates following *Gpc6* loss. Control cells exhibited progressive matrix mineralisation throughout the differentiation time course (Figure 6A). In contrast, *Gpc6*-deficient cells displayed a marked delay in mineralisation, achieving levels comparable to those of controls approximately 10 days later (Figure 6B). These findings indicate that loss of *Gpc6* delays progression through the hypertrophic differentiation programme, consistent with the delayed hypertrophic differentiation observed in *Gpc6*-deficient growth plates *in vivo*. To determine whether this delay was mediated by ERK1/2 signalling, as reported for PTPN11/SHP2-associated differentiation defects in which MEK inhibition alleviates delayed maturation,^66^ differentiating cultures were treated with low-dose trametinib to attenuate MEK–ERK1/2 activity. Trametinib treatment partially but reproducibly accelerated mineralisation in *Gpc6*-deficient cultures (Figure 6C). These findings indicate that ERK1/2 hyperactivation contributes to the delayed chondrocyte differentiation phenotype caused by *Gpc6* loss.

**Figure 6.**
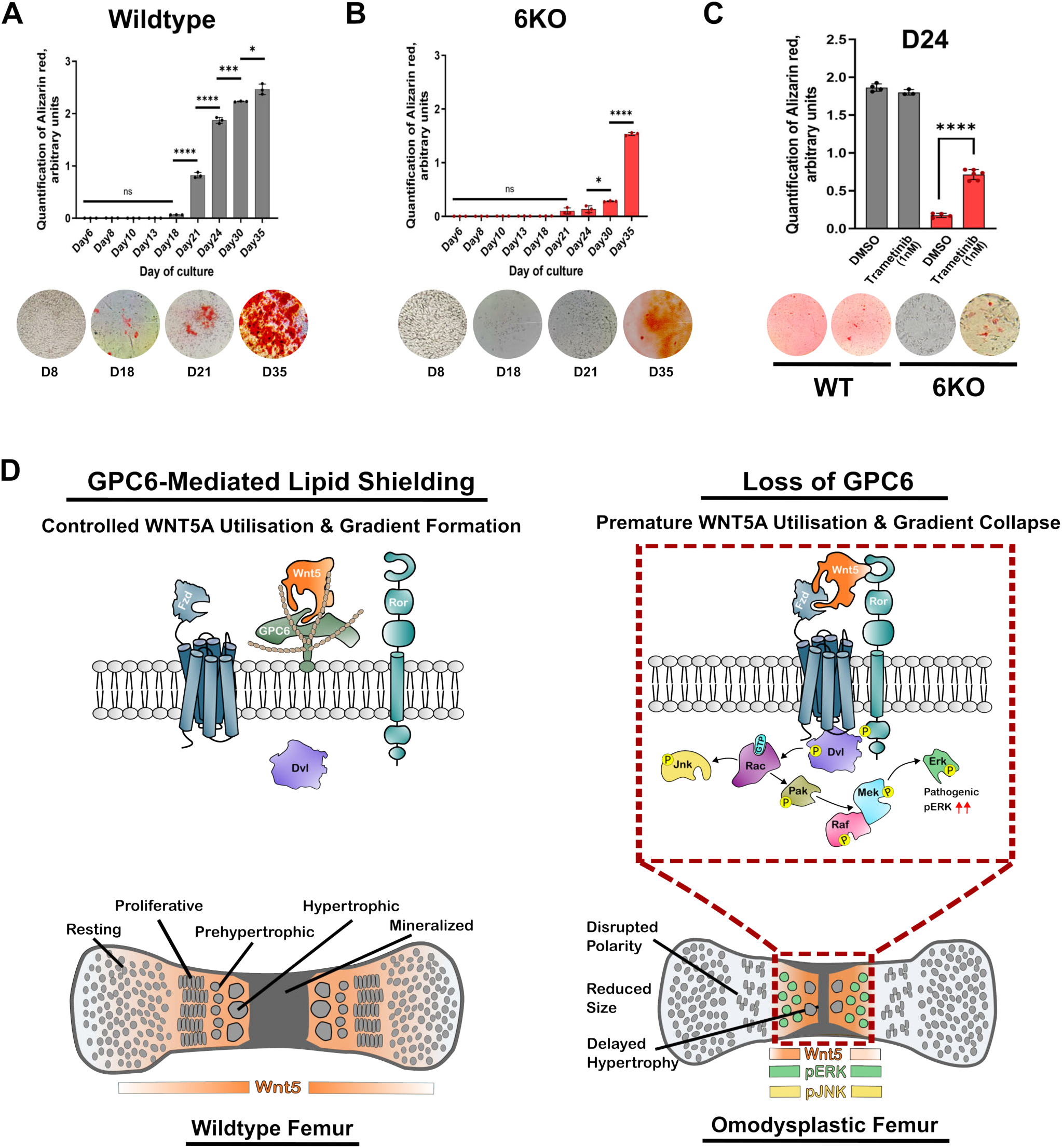
Aberrant ERK Signalling Links GPC6 Loss to Impaired Chondrocyte Differentiation. **(A)** Quantification of Alizarin Red S staining in wildtype (WT) ATDC5 cells over a 35-day differentiation time course across 9 time-points *(top)*, with corresponding representative staining images at days 8, 18, 21, and 35 of culture *(bottom)*. Bar graph data represent mean (*n* = 3) ± SD. Statistical significance between consecutive time-points was determined using unpaired t-test with Welch’s correction (\**P* = 0.0165, \*\*\**P* = 0.0004, \*\*\*\**P* < 0.00001\); ns: not significant). **(B)** Quantification of Alizarin Red S staining in Gpc6-deficient (6KO) ATDC5 cells over a 35-day differentiation time course across 9 time-points *(top)*, with corresponding representative staining images at days 8, 18, 21, and 35 of culture *(bottom)*. Bar graph data represent mean (*n* = 3) ± SD. Statistical significance between consecutive time-points was determined using unpaired t-test with Welch’s correction (\**P* = 0.0164, \*\*\*\**P* < 0.00001\); ns: not significant). **(C)** Quantification of matrix mineralisation in WT + DMSO (*n* = 4), WT + trametinib (*n* = 3), 6KO + DMSO (*n* = 4), and 6KO + trametinib (*n* = 6) cultures *(top)*, with corresponding representative staining images at days 24 of culture *(bottom)*. Attenuation of MEK-ERK signalling results in a partial restoration of mineralisation in Gpc6-deficient cells. Statistical significance was determined using an unpaired t-test with Welch’s correction (\*\*\*\**P* < 0.00001). **(D)** Model illustrating the proposed role of GPC6 in regulating the spatial utilisation of signalling-competent WNT5A. GPC6 binds lipid-modified WNT5A through a conserved lipid-binding pocket, transiently buffering signalling-competent ligand and limiting premature productive receptor engagement within the WNT5A-producing environment. This promotes ligand dispersal and tissue-scale gradient formation. Loss of GPC6-mediated lipid shielding increases local WNT5A utilisation, resulting in source-proximal activation of a non-canonical WNT-RAC-PAK-MEK-ERK signalling axis and collapse of the physiological WNT5A gradient. Together, these signalling alterations disrupt growth plate organisation, delay hypertrophic differentiation, and contribute to the skeletal abnormalities associated with GPC6 deficiency and Omodysplasia.

Together, these findings establish a functional consequence of ligand-dependent non-canonical WNT hyperactivation following *Gpc6* loss, linking activation of the RAC–PAK– MEK–ERK signalling axis to delayed hypertrophic chondrocyte differentiation. Thus, loss of GPC6 disrupts multiple coordinated processes required for normal growth plate development, including chondrocyte polarity, proliferation and timely hypertrophic differentiation. Our findings support a model in which GPC6 binds the lipid-modified WNT5A ligand through its conserved lipid-binding pocket, transiently shielding the ligand from premature productive engagement with Frizzled receptors within the WNT5A-producing environment (Figure 6D, left). Loss of GPC6-mediated lipid shielding therefore increases the availability of WNT5A for local receptor engagement, resulting in enhanced non-canonical WNT signalling and activation of the RAC–PAK–MEK–ERK cascade (Figure 6D, right). We propose that GPC6 therefore acts not simply to maintain WNT5A in a soluble, dispersible state, but to transiently buffer signalling-competent ligand and regulate its accessibility to receptors at the site of production; loss of this local buffering drives a RASopathy-like developmental pathology.

## Discussion

Morphogen gradients coordinate tissue patterning, morphogenesis and organ development by controlling where and when signalling occurs. Our findings show that gradient formation depends not only on mechanisms that preserve the extracellular mobility of signalling-competent ligand, but also on mechanisms that constrain where that ligand is utilised. By selectively disrupting a conserved WNT lipid-binding interface in GPC6, we genetically separate glypican-mediated lipid engagement from other glypican functions *in vivo*. GPC6-mediated WNT lipid shielding is essential for normal skeletal development and restricts source-proximal utilisation of signalling-competent WNT5A, allowing ligand dispersal before productive receptor engagement and thereby promoting tissue-scale signalling.

Our structure-guided Gpc6^A116W^ allele provides a tractable model for a separation-of-function analysis of WNT lipid shielding. It disrupts the predicted WNT lipid-binding interface while preserving GPC6 expression, maturation and detectable heparan sulfate modification.

Gpc6^A116W^ recapitulates the major skeletal, craniofacial and signalling abnormalities of complete Gpc6 loss despite retaining heparan sulfate chains. However, the skeletal and craniofacial abnormalities are less severe than those of the null, and cleft palate is not observed, indicating that the A116W allele does not fully phenocopy complete GPC6 deficiency. This incomplete phenotypic overlap suggests that some GPC6 functions are retained despite disruption of the lipid-binding interface. One possibility is that A116W does not completely eliminate WNT lipid binding *in vivo*. Although A116W abolishes detectable binding to the lipidated WNT peptide in our biochemical assays, interactions between full-length GPC6 and WNT5A could provide additional protein–protein contacts that stabilise residual lipid-dependent binding *in vivo*. Alternatively, some developmental functions of GPC6 may be mediated independently of direct WNT lipid binding. In particular, its HS-GAG chains can mediate interactions with WNTs and other growth factors, providing a potential mechanism for residual GPC6 function in tissues in which the A116W allele produces milder phenotypes than complete Gpc6 loss. Thus, the milder A116W phenotype may reflect residual WNT lipid engagement, additional HS-GAG-mediated interactions, or both. These findings establish WNT lipid engagement as an important function of GPC6 during skeletal development and provide a genetic framework for separating lipid binding from other glypican activities *in vivo*.

Unexpectedly, disruption of GPC6-mediated lipid engagement does not simply reduce WNT signalling but redistributes it within the tissue. Loss of GPC6 disrupts WNT5A gradient architecture, reducing ligand distribution across the developing growth plate while concentrating signalling activity within the WNT5A-producing prehypertrophic domain. Notably, signalling increases at the source despite reduced WNT5A abundance, demonstrating that reduced morphogen abundance does not necessarily indicate reduced pathway activity. This spatial redistribution provides a framework for understanding the apparently opposing consequences of *Gpc6* loss: increased WNT5A utilisation near the source produces a local gain of signalling, whereas depletion of the ligand available for more distal regions results in a corresponding loss of tissue-scale WNT5A signalling. These findings suggest that reduced proliferation, disrupted columnar organisation and impaired maturation of the growth plate are likely to reflect, at least in part, insufficient WNT5A signalling away from the source, occurring alongside excessive signalling within the ligand-producing domain. GPC6 lipid binding therefore regulates the allocation of a finite pool of signalling-competent WNT5A between different regions of the tissue, rather than simply determining the overall amount of signalling. This redistribution may also explain why previous skeletal analyses failed to detect non-canonical WNT signalling defects following *Gpc6* loss.^33^ Bulk measurements can obscure opposing changes in pathway activity across distinct tissue domains, emphasising the importance of spatially resolved analysis when interpreting morphogen signalling.

The increased source-proximal signalling with reduced WNT5A abundance following *Gpc6* loss is consistent with signalling-competent WNT5A being utilised by Frizzled receptors near the site of production, providing an alternative means of shielding the lipidated ligand in the absence of GPC6, albeit one that consequently drives excess signalling. These findings suggest that source-proximal WNT signalling is not normally restrained by the absence of receptive signalling machinery, but by mechanisms that regulate ligand access to available receptors. Extracellular buffering may therefore restrain autocrine signalling without rendering WNT-producing cells refractory to WNT, preserving their capacity to respond should signalling requirements change during development or tissue repair.

A related question is how WNT5A transitions from a GPC6-associated extracellular state to productive Frizzled engagement. Because both GPC6 and Frizzled interact with the lipidated region of WNT5A, signalling may require a transition between protected and receptor-accessible states. This possibility places our findings in the context of current models of extracellular WNT transport. A recent model proposed that newly secreted WNT is transferred from Wntless (WLS), which binds the lipid modification of WNT within a hydrophobic pocket and delivers WNT to the cell surface for secretion,^74–77^ to soluble carriers such as sFRPs and WIF1 and subsequently to cell-surface glypicans in receiving cells, where glypican interactions may facilitate ligand release for Frizzled engagement and signal transduction.^78^ However, this model does not yet explain how WNT is handed between these extracellular partners or how premature receptor engagement is avoided during transit. Our findings suggest that this sequence may capture a later stage of WNT trafficking, while glypicans also act earlier in the extracellular journey of newly secreted ligand. Rather than functioning solely at receiving cells to promote ligand release to Frizzled, GPC6 transiently shields newly secreted WNT before its subsequent loading into dispersal pathways such as soluble carriers, extracellular vesicles or cytonemes. Our findings do not exclude roles for sFRPs, WIF1 or other extracellular carriers in downstream ligand transfer and dissemination, but suggest that glypican-mediated lipid shielding may precede these interactions and provide a protected interval between WNT secretion and productive receptor engagement.

GPC6 may thus regulate not simply how WNT is transported, but when signalling-competent ligand becomes accessible to its receptors.

Consistent with this model, WNT5A gradient collapse generates a source-confined signalling state that engages RAC–PAK–MEK–ERK within the WNT5A-producing compartment. Thus, defective morphogen organisation can do more than reduce long-range signalling: altered extracellular ligand distribution can be propagated into aberrant intracellular signalling within the ligand-producing domain. RAF and MEK inhibition abolished ERK activation following *Gpc6* loss, indicating that local non-canonical WNT hyperactivation is channelled through the canonical RAF–MEK–ERK cascade. PAK provides a mechanistic link between these pathways: PAK1 can phosphorylate MEK1 at Ser298, which promotes efficient RAF-dependent MEK activation and enhances functional coupling of MEK1 to ERK.^53,54^ Consistent with this mechanism, Gpc6 loss increased MEK1 Ser298 phosphorylation, while PAK inhibition reduced both Ser298 phosphorylation and ERK activation. Together, these findings identify RAC–PAK–dependent potentiation of the RAF–MEK–ERK cascade as a route through which altered extracellular WNT organisation can generate pathological ERK1/2 activation.

The convergence of extracellular WNT5A dysregulation and intracellular ERK1/2 activation provides a mechanistic explanation for the phenotypic relationships between GPC6-associated omodysplasia, Robinow syndrome and RASopathies. Although GPC6-associated skeletal defects have largely been interpreted through altered Hedgehog signalling,^33^ our findings identify disruption of WNT5A organisation as a previously unrecognised pathogenic mechanism and provide a molecular explanation for the substantial overlap between omodysplasia and Robinow syndrome.^30,32^ However, GPC6 deficiency differs fundamentally from the loss of WNT5A pathway activity caused by mutations in core pathway components such as *WNT5A*, *ROR2* or *DVL*. Rather than uniformly reducing WNT5A signalling, loss of GPC6 redistributes signalling within the tissue, reducing distal signalling in the developing growth plate while simultaneously increasing signalling within the ligand-producing domain. This spatially opposing signalling state — loss of tissue-scale WNT5A signalling alongside source-confined hyperactivation — directly engages the RAC–PAK–MEK–ERK cascade.

Strikingly, this generates a RASopathy-like developmental signalling state without mutation or direct perturbation of the intracellular MAPK machinery. Thus, disruption of extracellular WNT organisation can produce pathological RAS–ERK activation through altered ligand utilisation, revealing an extracellular route to a signalling state otherwise associated with lesions in intracellular pathway components.

ERK1/2 hyperactivation is unlikely to account for the entirety of the *Gpc6* phenotype. We favour a model in which gradient collapse has dual pathological consequences: loss of tissue-scale WNT5A signalling disrupts growth plate organisation and columnar polarity, whereas increased signalling within the ligand-producing domain drives RAC–PAK–MEK– ERK activation and delays chondrocyte hypertrophic differentiation. GPC6-associated omodysplasia therefore combines impaired tissue-scale WNT5A signalling, reminiscent of Robinow syndrome, with a local gain-of-function RAS–ERK state characteristic of RASopathies,^79,80^ defining omodysplasia as a molecular hybrid with features of both developmental disease states.

Collectively, our findings establish extracellular morphogen organisation as a fundamental layer of developmental signalling regulation. Glypicans regulate not only where signalling-competent ligands are distributed, but where they become available for productive receptor engagement. By restricting immediate source-proximal utilisation of signalling-competent WNT5A, glypican-mediated lipid shielding coordinates extracellular ligand availability with tissue-scale morphogen dispersal. In this view, lipid shielding controls the spatial allocation of a finite pool of signalling-competent ligand: loss of shielding increases WNT5A utilisation at the source while depleting ligand available to more distant cells, thereby coupling local signalling gain to tissue-scale signalling loss. More broadly, this work provides a framework for understanding how extracellular mechanisms determine where, when and by which cells signalling occurs, and suggests that developmental disease may arise not only through mutations affecting intracellular signalling pathways, but also through disruption of extracellular morphogen organisation.

### Limitations of the study

Our findings establish GPC6-mediated lipid shielding as a regulator of WNT5A spatial utilisation during skeletal development, but the generality of this mechanism remains to be established. GPC6 expression overlaps with WNT5A in multiple embryonic tissues, raising the possibility that glypican-mediated lipid shielding contributes to WNT5A signalling architecture beyond the developing skeleton and that analogous mechanisms may operate between other glypican–WNT pairs. Although our cellular studies establish that WNT5A/B is required for the increased ERK signalling observed following loss of GPC6, we have not genetically tested whether WNT5A/B is similarly required for the elevated ERK activity *in vivo*. The spatial correspondence between the GPC6 and WNT5A phenotypes, together with the collapse of the WNT5A signalling range in *Gpc6*-deficient skeletal tissue, supports WNT5A as the principal ligand underlying this response; however, contributions from other WNT ligands, including the partially redundant WNT5B, cannot be excluded. This uncertainty does not bear on our central conclusion, which concerns the spatial regulation of ligand access by GPC6 rather than the identity of the specific WNT ligand involved. In addition, although our data strongly support a shift towards source-proximal Frizzled utilisation following loss of GPC6, whether loss of GPC6 also causes additional ligand loss through extracellular destabilisation remains unresolved. More broadly, the molecular mechanisms underpinning the handoff of WNT5A between GPC6, Frizzled and other extracellular WNT-binding or dispersal systems remain to be directly defined. Determining how these extracellular interactions coordinate ligand shielding, dispersal and receptor engagement will be an important direction for future studies.

## Methods

### Structural modelling of GPC6 in a lipid-bound conformation

The mouse GPC6 amino-acid sequence (UniProt Q9R087) was submitted to the AlphaFold 3 server (https://alphafoldserver.com) for structure prediction. The resulting model adopted the characteristic glypican fold and closely resembled the apo crystal structure of *Drosophila* Dally-like protein (Dlp; PDB 3ODN), including conservation of the C-lobe α-helical architecture surrounding the putative lipid-binding cavity.

To model GPC6 in a lipid-bound conformation, a template-based model was subsequently generated using SWISS-MODEL (https://swissmodel.expasy.org) with the lipid-bound Dlp structure (PDB 6XTZ) as the structural template. The resulting model was used to assess conservation of the hydrophobic cavity, positioning of the palmitoleoylated serine within the predicted GPC6 lipid-binding pocket, and the structural consequences of substitutions at the cavity entrance.

### Recombinant GPC6 protein expression and purification

cDNAs encoding the human GPC6 (UniProt Q9Y625) core ectodomain (residues Asp24– Thr488), either wild type or carrying the A116W substitution, were cloned into the mammalian expression vector pHLsec (Addgene #99845), which introduces a C-terminal 6×His tag.

Recombinant GPC6core wild-type and GPC6core A116W proteins were expressed by transient transfection of HEK293F suspension cells maintained in Expi293 Expression Medium (ThermoFisher) at 37°C with 8% CO_2_. For each litre of cell culture, 0.5 mg plasmid DNA and 1 mg polyethylenimine (PEI) were separately diluted in 25 ml FreeStyle 293 Expression Medium (ThermoFisher). The diluted DNA and PEI solutions were mixed and incubated for 10 min at room temperature before being added directly to the suspension culture. Conditioned medium containing secreted recombinant proteins were harvested 4 days post-transfection and buffer exchanged into phosphate buffer (25 mM phosphate, pH 8.0, 250 mM NaCl) using a QuixStand benchtop tangential flow filtration system (GE Healthcare) fitted with a 60-cm Xampler cartridge (GE Healthcare) with a 10-kDa nominal molecular weight cut-off membrane. The buffer-exchanged medium was applied to a 5 ml HisTrap Excel affinity column (Cytiva). Following washing with phosphate buffer containing 5 mM imidazole, bound protein was eluted with phosphate buffer supplemented with 300 mM imidazole. Eluted protein was further purified by size-exclusion chromatography on a Superose 6 Increase 10/300 column (Cytiva) equilibrated in 10 mM HEPES (pH 7.4) and 150 mM NaCl. Fractions containing purified GPC6core were pooled for subsequent biochemical analyses.

### Lipidated WNT7A peptide synthesis

A lipidated peptide corresponding to residues 204–213 of mouse WNT7A was synthesised as previously described.^18^ The peptide contained a palmitoleoylated serine modification (S206 C16:1 ox) together with a C-terminal biotin modification to enable detection through fluorescent streptavidin binding. The peptide sequence was: NH₂-K(Cys-Ser-)HGV(Ser-palmitoleoyl)GS(Cys-Ser-)TTKT-PEG-Bio. The expected molecular mass of the synthetic peptide (1970.5 Da) was confirmed by mass spectrometry.

For fluorescence polarisation experiments, fluorescent WNT7A lipid peptide (Wnt7a_lip_5-FAM) and the corresponding non-lipidated control peptide were synthesised as previously described,^18^ except that 5-FAM was coupled to a Universal-PEG resin.

### Fluorescence polarisation

Fluorescence polarisation (FP) measurements were performed using a JASCO FP-8500 spectrofluorometer equipped with excitation and emission polarisers using a 3 × 3 mm quartz cuvette. Fluorescent ligands (50 nM) were prepared in assay buffer containing 25 mM HEPES (pH 7.5) and 200 mM NaCl, and measurements were performed at 20°C. Increasing concentrations of purified recombinant GPC6^core^ were added sequentially to the fluorescent ligand, and fluorescence anisotropy was recorded following each addition. Excitation and emission wavelengths and slit widths were selected according to the fluorophore used (5-FAM: excitation 490 nm, emission 530 nm; excitation slit 5 nm, emission slit 10 nm).

Fluorescence anisotropy data were fitted by nonlinear regression using a quadratic binding equation to account for ligand depletion. The equilibrium dissociation constant (K_D_), together with the anisotropy values of the free ligand (A_min_) and protein-bound ligand (A_max_), were determined by fitting the experimental data while fixing the fluorescent ligand concentration.

### Protein–lipid binding by fluorescence-detection size-exclusion chromatography (FSEC)

Binding of GPC6 to the lipidated WNT7A-derived peptide was assessed by fluorescence-detection size-exclusion chromatography (FSEC). The palmitoleoylated, biotinylated WNT7A peptide was mixed with streptavidin–Alexa Fluor 488 (Invitrogen) to generate a fluorescently detectable peptide complex. The labelled peptide was subsequently mixed with purified wild-type or GPC6core or GPC6core^A116W^ protein. Protein–peptide mixtures were incubated for 1 hr at 4°C in binding buffer consisting of 10 mM HEPES (pH 7.4), 150 mM NaCl and 0.01% Tween-20. GPC6^core^ proteins were used at a final concentration of 2 μM with approximately 20 μM lipidated peptide and 20 μM streptavidin–Alexa Fluor 488. Following incubation at room temperature for 1 hour, samples were analysed by fluorescence-detection size-exclusion chromatography using a Superose 6 Increase 3.2/300 column (Cytiva) connected to a Shimadzu Prominence HPLC system equipped with an RF-10AXL fluorescence detector. The running buffer was identical to the incubation buffer above. Chromatography was performed at a flow rate of 0.08 ml min⁻¹. Fluorescence was monitored using excitation at 505 nm and emission at 512 nm. UV absorbance at 280 nm was recorded simultaneously to compare protein elution profiles with fluorescence signals. A control sample containing the lipidated peptide–streptavidin–Alexa Fluor 488 complex in the absence of GPC6 was analysed under identical conditions. Association of the fluorescent peptide with GPC6 was assessed by its shift to an earlier elution volume coincident with the protein-containing fractions.

### Mouse husbandry and allele generation

The *Gpc6*^tm2b(EUCOMM)Wtsi^ GPC6 knockout (*Gpc6*^tm2b^) mouse line was generated as described previously.^41^ Briefly, the *Gpc6*^tm2a(EUCOMM)Wtsi^ mouse strain (MGI:4453713), generated by the Wellcome Trust Sanger Institute and obtained from the Infrafrontier repository, was used as the starting line. The *Gpc6*^tm2a(EUCOMM)Wtsi^ allele was converted to the constitutive null *Gpc6*^tm2b(EUCOMM)Wtsi^ allele by Cre-mediated recombination.

The *Gpc6*^A116W^ and *Gpc6*^-10bp^ alleles were generated by CRISPR/Cas12a-mediated genome editing using homology-directed repair in mouse zygotes. Zygotes obtained from superovulated C57BL/6 females were incubated with single-stranded adeno-associated virus serotype 6 (ssAAV6; 2 × 10^11^ genome copies ml^−1^) carrying the donor DNA template encoding the A116W substitution. Following delivery of the repair template, zygotes were electroporated with Acidaminococcus sp. Cas12a (AsCas12a) protein (200 ng μl^−1^; Integrated DNA Technologies) and Cas12a guide RNA (200 ng μl^−1^; Integrated DNA Technologies) targeting the *Gpc6* locus using a NEPA21 electroporator (Sonidel, Japan).

Electroporation was performed using four 40 V pulses of 3.5 ms duration separated by 50 ms intervals. Embryos were cultured overnight in M16 medium to the two-cell stage before transfer into pseudopregnant CD-1 recipient females. Founder mice were genotyped by PCR, and the *Gpc6*^A116W^ allele was confirmed by Sanger sequencing. In addition to the repaired allele, embryos that failed to undergo homology-directed repair generated independent CRISPR-induced alleles containing frameshift mutations within *Gpc6*. One founder carrying a 10 bp deletion predicted to generate a null allele (*Gpc6*^-10bp^) was identified, confirmed by Sanger sequencing and maintained as an independent knockout line for phenotypic analyses. Routine colony genotyping was subsequently performed by Transnetyx (USA).

All animal procedures were approved by the Babraham Institute Animal Welfare and Ethical Review Body. Animal husbandry and experimentation were conducted in accordance with European Union and United Kingdom Home Office legislation and institutional guidelines under project licence PP4567823. Mice were bred and maintained in the Babraham Institute Biological Support Unit. Timed matings were established, and the day on which a copulation plug was observed was designated embryonic day (E)0.5. Animals were maintained at an ambient temperature of approximately 19–21°C and relative humidity of 52% under a 12 h light/12 h dark cycle incorporating 15 min dawn and dusk transition periods.

### Whole-mount skeletal preparations

Whole-mount skeletal preparations were performed as previously described.^81^ Briefly, embryos were collected, eviscerated and fixed in 95% ethanol. For analysis of the appendicular skeleton, samples were stained overnight with 0.03% (w/v) Alcian Blue 8GX (Sigma) prepared in 70% ethanol containing 20% glacial acetic acid to visualise cartilage. Following extensive washing in 95% ethanol, specimens were cleared in 1% potassium hydroxide (KOH) and stained overnight with 0.005% (w/v) Alizarin Red S (Sigma) prepared in 1% KOH to visualise mineralised bone. Samples were subsequently cleared through increasing concentrations of glycerol in KOH before storage in 100% glycerol until imaging. For craniofacial skeletal analysis, skulls were dissected from fixed embryos, carefully cleared of surrounding soft tissue and processed separately. Isolated skulls were stained overnight with 0.005% (w/v) Alizarin Red S in 1% KOH, followed by clearing through glycerol/KOH solutions and storage in 100% glycerol before imaging.

### Micro-computed tomography

For craniofacial evaluation, dissected mouse heads were harvested from P1 mice. Specimens were fixed in 4% paraformaldehyde, skin was removed and heads were treated with 2% Lugol’s solution for 24 hrs and washed in PBS before imaging. High-resolution ex vivo micro-computed tomography (μCT) imaging was performed using a Nikon Metrology XT H 225 μCT system running XT 6.8 software and equipped with a Reflection 225 X-ray source. The X-ray source settings were operated at an energy voltage of 110 kV and an intensity current of 64 μA using the auto-defocus mode. For each scan, 1,080 angular projection images were acquired over a 360° rotation with an exposure time of 708 ms per frame. Scans were performed without additional filtration at an isotropic voxel size of 8.03 μm. Three-dimensional reconstructions were generated from raw projection datasets using Nikon CTPro software (V6.8). Regions of interest (ROIs) for craniofacial structures were segmented from the surrounding background using a fixed global threshold.

### Bone collection, fixation and cryosectioning

Embryonic mice were euthanised by hypothermia. Postnatal mice were euthanised either by carbon dioxide inhalation or cervical dislocation in accordance with institutional guidelines and UK Home Office regulations. Hindlimbs were dissected in ice-cold PBS, carefully cleared of skin and surrounding soft tissue and fixed in 4% paraformaldehyde for 48 hr at 4°C. Samples were processed without decalcification. Following fixation, tissues were washed extensively in PBS before sequential cryoprotection in 15% and 30% sucrose prepared in PBS at 4°C until the samples had fully equilibrated. Cryoprotected limbs were embedded in Optimal Cutting Temperature (OCT) compound (Tissue-Tek) with the sagittal plane oriented towards the cutting surface. Embedded samples were rapidly frozen by immersion in isopentane cooled on dry ice. Serial cryosections (6-8 μm for immunostaining and 8 μm for H&E staining) were collected using a Leica cryostat onto Superfrost microscope slides, air dried for at least 30 min and stored at −80°C until required. Immediately before histological or immunofluorescence staining, slides were brought to room temperature in PBS and residual OCT compound was removed by washing in PBS.

### Immunofluorescence staining and microscopy

Cryosections were permeabilised and blocked in PBSTx before incubation with primary antibodies diluted in blocking buffer for overnight at 4°C. Following washes in PBSTx, sections were incubated for 1 h at room temperature with Alexa Fluor 488-, Alexa Fluor 555- and/or Alexa Fluor 647-conjugated secondary antibodies (Molecular Probes; 1:2000) together with DAPI for nuclear counterstaining. Following further washes in PBSTx, slides were mounted using Mowiol (2.4% Mowiol 4-88 [poly(vinyl alcohol)], Sigma). Images were acquired using a Nikon Ti2 inverted fluorescence microscope equipped with 4×, 10×, 20×, 40×, 60× and 100× objectives, together with a Hamamatsu Flash4.0 sCMOS camera and a Nikon Ri2 colour camera. The primary antibodies used in immunofluorescence staining, along with their catalogue numbers, experimental purposes, and dilutions, are detailed in Table S1.

Image analysis included: Automated cell segmentation and planar cell polarity (PCP) orientation were analysed using the PCP Auto Count (PCPA) plugin in Fiji/ImageJ. Images were thresholded into binary masks to identify cell boundaries and localised polarity signals. The PCPA plugin calculated individual cell vectors and mean orientation angles relative to a 0° anatomical baseline. Tracking accuracy was verified manually before data aggregation. Angular frequencies were then plotted as circular histograms (rose diagrams) using the built-in PCPA utility. For orientation analysis, a minimum of 494 wild-type cells, 507 point-mutant cells, and 829 knockout cells across 13 independent regions were quantified. To evaluate morphological changes, individual cell aspect ratios were calculated using the standard shape descriptors utility in Fiji/ImageJ. This morphological quantification was performed on a subset of 205 wild-type cells, 211 point-mutant (A116W) cells, and 211 knockout cells. The angular orientation distributions of these specific subsets were subsequently visualised as half-rose plots generated in RStudio (version 2026.07.1+147).

Relative fluorescence intensity was quantified using Fiji/ImageJ software. Multichannel confocal images were separated into individual channels to isolate target signals from the DAPI nuclear control. For each channel, background fluorescence was subtracted from the raw mean gray values of the regions of interest (ROIs). To account for variations in sample preparation and cell density, the background-corrected intensity of each target channel was normalized to the corresponding DAPI signal intensity. Final quantitative data are expressed as the relative fluorescence intensity ratio (Target/DAPI).

To quantify chondrocyte proliferation, fluorescence images of growth plate sections were analyzed using Fiji/ImageJ software. The total cell population within a standardized field of view was quantified via automated segmentation of DAPI-stained nuclei. Mitotically active chondrocytes were identified by positive immunoreactivity for phospho-Histone H3 (pHH3^+^). The proliferation index was calculated as the percentage of pHH3^+^ cells relative to the total number of DAPI-stained nuclei. Image processing, thresholding, and automated particle counting were executed using a custom macro to ensure reproducibility.

### Embryonic femur explant culture

Embryonic femur explants were cultured as previously described.^82^ Briefly, E14.5 femurs were dissected from *Gpc6*^tm2b^ embryos and wild-type littermates, carefully cleared of surrounding soft tissue in ice-cold PBS and cultured individually in 24-well plates at 37°C in a humidified atmosphere containing 5% CO₂. Explants were maintained in serum-free DMEM (Gibco) supplemented with 0.2% bovine serum albumin, 0.5 mM L-glutamine, 40 U ml⁻¹ penicillin–streptomycin (Gibco), 0.05 mg ml⁻¹ ascorbic acid (Sigma) and 1 mM β-glycerophosphate (Sigma). Where indicated, contralateral femurs from the same embryo were cultured for 24 h in the presence of either vehicle (DMSO) or the Porcupine inhibitor C59 (50nM). Following treatment, explants were fixed in 4% paraformaldehyde and processed for immunofluorescence.

### Cell culture and Transfection

Murine ATDC5 chondrogenic cells (ECACC, Cat. No. 99072806) were maintained in DMEM/F-12 (1:1) supplemented with GlutaMAX™ I and 10% fetal bovine serum. For chondrogenic differentiation, cells were seeded at a density of 6,000 cells cm⁻² in 12-well plates and cultured in differentiation medium consisting of DMEM/F-12 (1:1) supplemented with GlutaMAX™ I, 5% fetal bovine serum, 1% insulin–transferrin–selenium (ITS), 1% sodium pyruvate and 0.5% gentamicin (Invitrogen), as previously described.^46^ Cells were cultured for six days until confluence before differentiation medium was supplemented with 10 mM β-glycerophosphate and 50 μg ml⁻¹ L-ascorbic acid 2-phosphate. Cultures were maintained for up to 35 days at 37°C in a humidified atmosphere containing 5% CO₂, with medium replaced every 2–3 days. ATDC5 cells were transiently transfected using Lipofectamine™ 3000 (Thermo Fisher Scientific) according to the manufacturer’s instructions.

NIH 3T3 fibroblasts (ATCC, Manassas, VA, USA) were routinely maintained in Dulbecco’s modified Eagle medium (DMEM) supplemented with 2 mM L-glutamine, 100 U ml⁻¹ penicillin, 100 μg ml⁻¹ streptomycin (Sigma-Aldrich) and 10% newborn calf serum.

Drosophila Schneider 2 (S2) cells were obtained from the Drosophila Genomics Resource Center (DGRC) and maintained at 25°C in Schneider’s medium (Sigma) supplemented with 10% fetal bovine serum (Life Technologies), L-glutamine and 0.1 mg ml⁻¹ penicillin– streptomycin.

The plasmid encoding V5-tagged GPC6^ΔGPI^ (pMT-GPC6^ΔGPI^-V5) was generated as described previously.^18^ The A116W substitution was introduced by replacing the corresponding coding region with a synthetic DNA fragment (Integrated DNA Technologies) containing the desired mutation using standard restriction enzyme cloning. Plasmids were transiently transfected using Effectene Transfection Reagent (Qiagen) according to the manufacturer’s instructions. Conditioned medium was collected 24 h after transfection and concentrated using 10-kDa molecular weight cut-off Amicon Ultra centrifugal filters (Millipore). Cell lysates were prepared in parallel, and expression of GPC6 proteins in conditioned medium and cell lysates was analysed by immunoblotting using an anti-V5 antibody.

### CRISPR/Cas9-mediated genome editing

*Gpc6*- and *Gpc4*-deficient NIH 3T3 and ATDC5 cell lines were generated using CRISPR/Cas9-mediated genome editing. Single-guide RNAs were designed using the CRISPR Design Tool CHOPCHOP,^83^ cloned into pSpCas9(BB)-2A-Puro (Addgene plasmid #48139) and transfected using PolyFect transfection reagent (Qiagen). Following puromycin selection (Cambridge Bioscience), single-cell clones were isolated by limiting dilution in 96-well plates. Targeted editing was confirmed by Sanger sequencing and immunoblotting. sgRNA sequences and PCR primers used for clone validation are provided in the Table S2B.

### Pharmacological treatments

For acute signalling experiments in ATDC5 cells, cells were treated with the indicated inhibitors for 3 h before lysis. The following compounds were used: RAC inhibitor EHT1864 (MCE, 754240-09-0, 10 µM), PAK inhibitors FRAX597 (MCE, 1286739-19-2, 100 nM) and IPA-3 (MCE, 42521-82-4, 20 µM), MEK inhibitor trametinib (Selleck UK, 871700-17-3, 100 nM), pan-RAF inhibitor MLN2480 (Selleck UK, 1096708-71-2, 10 µM). To inhibit WNT secretion, cells were treated overnight with the Porcupine inhibitor C59 (MCE, 1243243-89-1, 10 nM) before analysis. Vehicle control cells received an equivalent concentration of DMSO.

### Western blotting

Cells were lysed in buffer containing 50 mM HEPES, 150 mM NaCl, 10% glycerol, 1% Nonidet P-40 and 1 mM EDTA supplemented with protease and phosphatase inhibitors (Sigma-Aldrich). Protein concentrations were determined using the Pierce™ BCA Protein Assay Kit (Thermo Fisher Scientific). Equal amounts of protein were resolved by SDS– PAGE and transferred onto Hybond-ECL membranes (Amersham). Membranes were blocked in 5% skimmed milk prepared in Tris-buffered saline containing 0.05% Tween-20 (TBST) before incubation with primary antibodies diluted in blocking buffer. Following washing in TBST, membranes were incubated with horseradish peroxidase-conjugated secondary antibodies and developed using enhanced chemiluminescence (GE Healthcare). The primary antibodies used in western blotting, along with their catalogue numbers, experimental purposes, and dilutions, are detailed in Table S1.

### RNA extraction, RT-PCR and quantitative PCR

Mouse tissues and cell pellets were harvested; homogenised and total RNA was extracted using the RNeasy Mini Kit according to the manufacturer’s instructions. First-strand cDNA was synthesised from total RNA using oligo(dT) primers and the Precision nanoScript2 Reverse Transcription Kit (Primer Design). Quantitative real-time PCR was performed using Maxima® SYBR Green qPCR Master Mix (Thermo Fisher Scientific) on a QuantStudio 5 Real-Time PCR System (Applied Biosystems). Crossing point (Cp) values were determined using QuantStudio 3/5 Real-Time PCR Software (Thermo Fisher). Relative gene expression was calculated using the 2^−ΔΔCt^ method with *Gapdh* as the reference gene. Primer sequences are provided in Table S2A.

### Alizarin Red S staining

Mineral deposition in differentiated ATDC5 cultures was assessed by Alizarin Red S staining as previously described.^46^ Briefly, cultures were fixed in 4% paraformaldehyde and stained with 2% Alizarin Red S solution (Sigma; pH 4.2) for 5 min at room temperature. Following staining, cultures were washed extensively with distilled water and imaged. For quantitative analysis of mineralisation, bound Alizarin Red S was extracted using 10% cetylpyridinium chloride for 10 min, and absorbance was measured at 570 nm using a microplate spectrophotometer.

### Statistical analysis

Statistical analyses were performed using GraphPad Prism 6 (La Jolla, CA, USA). Data are presented as mean ± s.d. unless otherwise indicated. Statistical tests were selected according to the experimental design and are specified in the corresponding figure legends. Comparisons between two groups were performed using an unpaired two-tailed Student’s *t*-test with Welch’s correction. Angular distributions were analysed using the Kolmogorov-Smirnov test. Multiple-group comparisons were performed using a one-way analysis of variance (ANOVA) followed by Dunnett’s post-hoc test as appropriate. The number of biological replicates (*n*), the definition of technical replicates, and the experimental unit are indicated in the corresponding figure legends. A *P*-value of *<0.05* was considered statistically significant.

## Author Contributions

Y.K. conceived and contributed to study design, designed and performed experiments, analysed data, and contributed to manuscript preparation. Y.Z. contributed to study design, performed structural modelling and biochemical experiments, including size-exclusion chromatography analyses, and contributed to manuscript preparation. A.X.X. performed experiments and analysed data. A.N. generated the Gpc6A116W and Gpc6-10bp mouse alleles using CRISPR/Cas12a-mediated genome editing. A.-M.A. performed experiments and analysed data, including fluorescence polarisation analyses. G.T. generated and provided the Gpc6tm2b mouse line, performed WNT5A staining of tibial tissue, and contributed to ex vivo explant studies. B.D.K. supervised G.T., provided mouse tissues, and contributed to ex vivo explant studies. E.Y.J. supervised Y.Z., contributed to study design, and provided intellectual input. I.J.M. conceived and supervised the study, designed experiments, analysed data, and wrote the manuscript with input from all authors.

## Use of Generative AI

Generative AI tools were used solely as writing and editing aids during the preparation of this manuscript. All experimental design, data collection, data analysis, interpretation and scientific conclusions were performed exclusively by the authors without AI assistance.

## Acknowledgements

This work was supported by a Sir Henry Dale Fellowship jointly funded by Wellcome and the Royal Society (221817/Z/20/Z), and by the Biotechnology and Biological Sciences Research Council (BBSRC) through the Babraham Institute Strategic Programme (BBS/E/B/000C0531), the Babraham Institute’s BBSRC Core Capability grant (CCG) (BB/CCG2210/1), and the Institute Development grant (BB/IDG2210/1). Bernard D. Keavney is supported by a British Heart Foundation Personal Chair. Gennadiy Tenin was supported by a British Heart Foundation Programme Grant (RG/F/21/110050). E. Yvonne Jones is supported by Wellcome Trust award (223133/Z/21/Z). We are grateful to Alberto Rosello-Diez for technical advice on bone explant culture conditions; Simon Walker and the Babraham Institute Imaging Facility for advice and assistance with imaging; the Babraham Institute Biological Support Unit, Stores, BICS, and Technical Services for support with mouse husbandry, experimental work, and research infrastructure. We thank Simon Cook and members of the Cook laboratory for technical advice on MAPK signalling, helpful discussions, and critical reading of the manuscript. We also thank Rahul Samant for critical reading of the manuscript, and Jean-Paul Vincent and David Willnow for insightful comments and suggestions. We thank all past and present members of the McGough laboratory for their support, discussions and contributions to this work. Finally, we thank the Babraham Institute Nursery for supporting our working families and contributing to an environment in which this research could be carried out.

---

**Figure S1.**
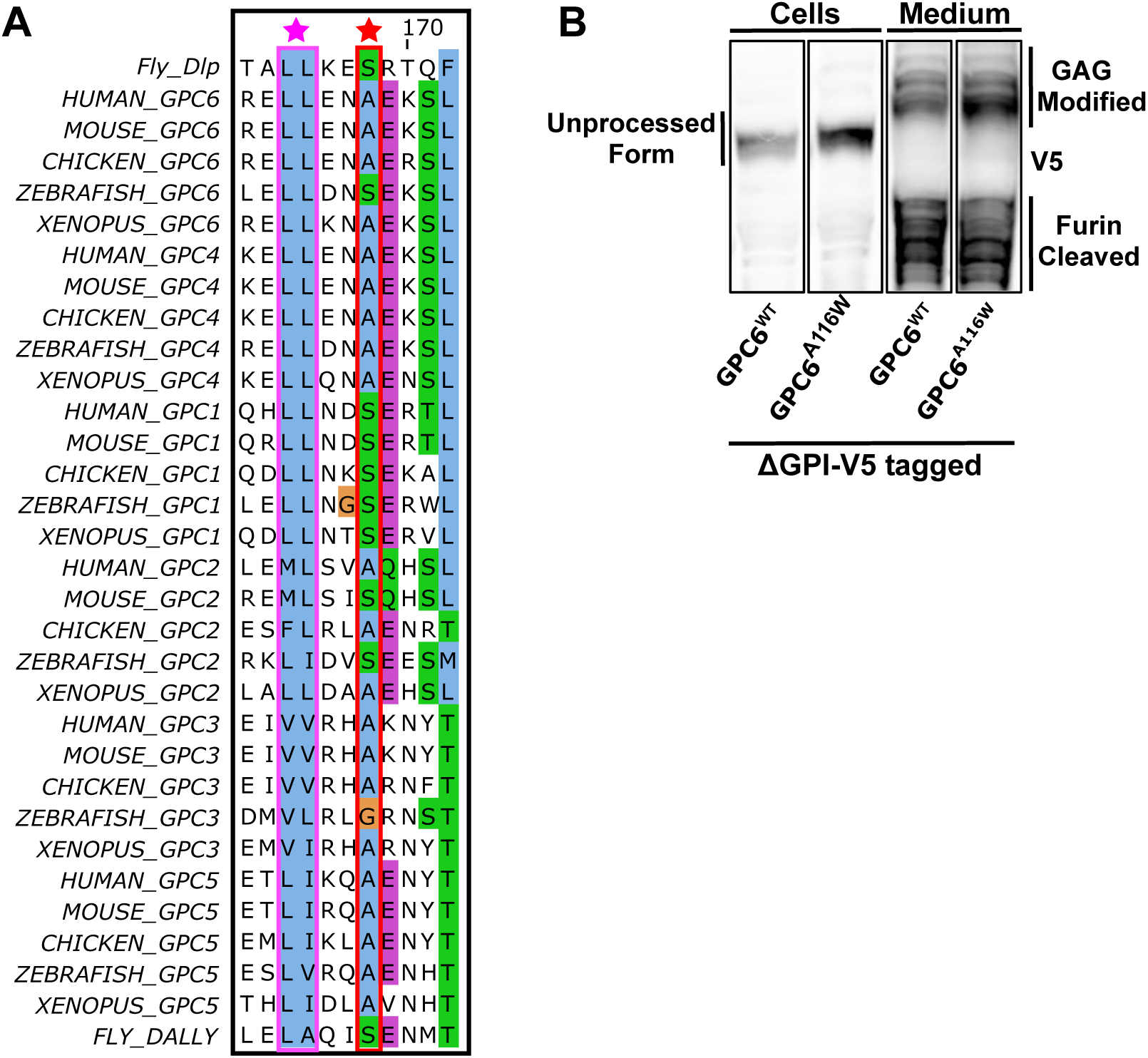
The GPC6 A116W Separation-of-Function Mutation Preserves Protein Processing and Secretion. **(A)** Multiple sequence alignment of glypican family members from human, mouse, chicken, zebrafish, *Xenopus*, and *Drosophila* highlighting conservation of residues surrounding the GPC6 WNT lipid-binding pocket. Conserved hydrophobic pocket residues (purple) and the conserved gatekeeper residue A116 (red) are indicated. **(B)** Immunoblot analysis of ΔGPI-V5-tagged WT and GPC6^A116W^ proteins in transfected cells and conditioned medium. The A116W mutation does not affect glycosaminoglycan (GAG) modification, furin processing, or secretion of GPC6.

**Figure S2.**
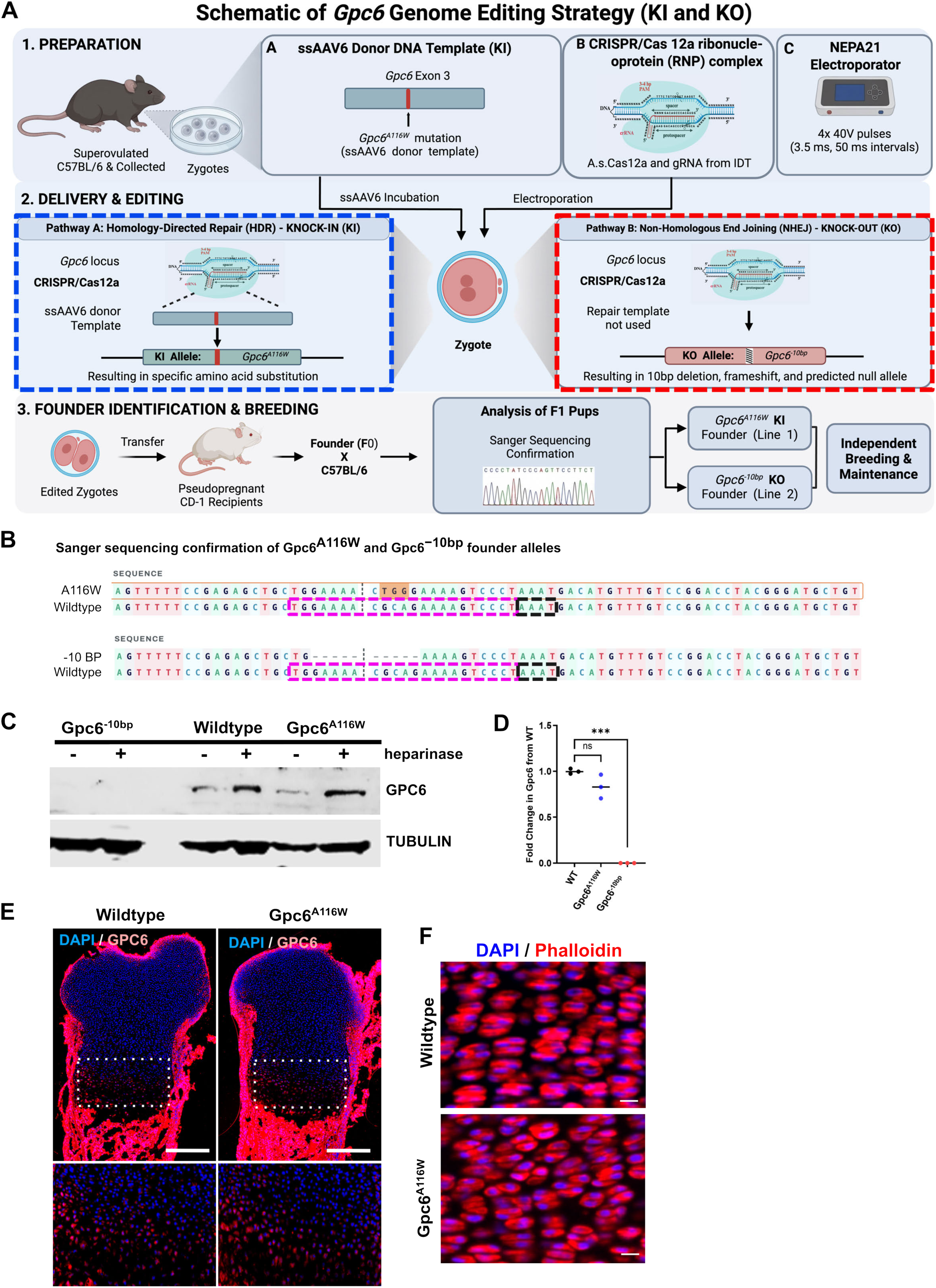
Generation and Validation of the *Gpc6^A116W^* and *Gpc6^−10bp^* Mouse Alleles. **(A)** Schematic of the CRISPR/Cas12a genome editing strategy used to generate the *Gpc6^A116W^* knock-in and *Gpc6^−10bp^* knockout alleles. **(B)** Sanger sequencing confirmation of the *Gpc6^A116W^* and *Gpc6^−10bp^* founder alleles. The Cas12a guide-target sequence is indicated by the purple dashed box and the PAM sequence by the black dashed box. The engineered A116W substitution and 10-bp deletion are indicated. **(C)** Immunoblot analysis of GPC6 protein in WT, *Gpc6^A116W^*, and *Gpc6^−10bp^*embryonic limb lysates treated with or without heparinase. **(D)** Quantitative RT-PCR analysis of *Gpc6* expression. *Gpc6* transcript levels were quantified in E13.5 limb extracts from WT, *Gpc6^A116W^* and *Gpc6^tm2b^*embryos and normalised to *Gapdh*. Data are expressed as mean fold induction relative to the WT control. Error bars indicate the standard deviation (SD) of five independent experiments. Statistical significance was determined using an unpaired *t*-test with Welch’s correction (\*\*\**P* = 0.0003). **(E)** Immunofluorescence analysis of GPC6 expression in E15.5 WT and *Gpc6^A116W^* femoral growth plates. Lower panels show higher-magnification views of the boxed regions. Scale bar 100 µm. **(F)** Phalloidin staining of columnar chondrocytes in P1 WT and *Gpc6^A116W^* femoral growth plates. Scale bar 100 µm.

**Figure S3.**
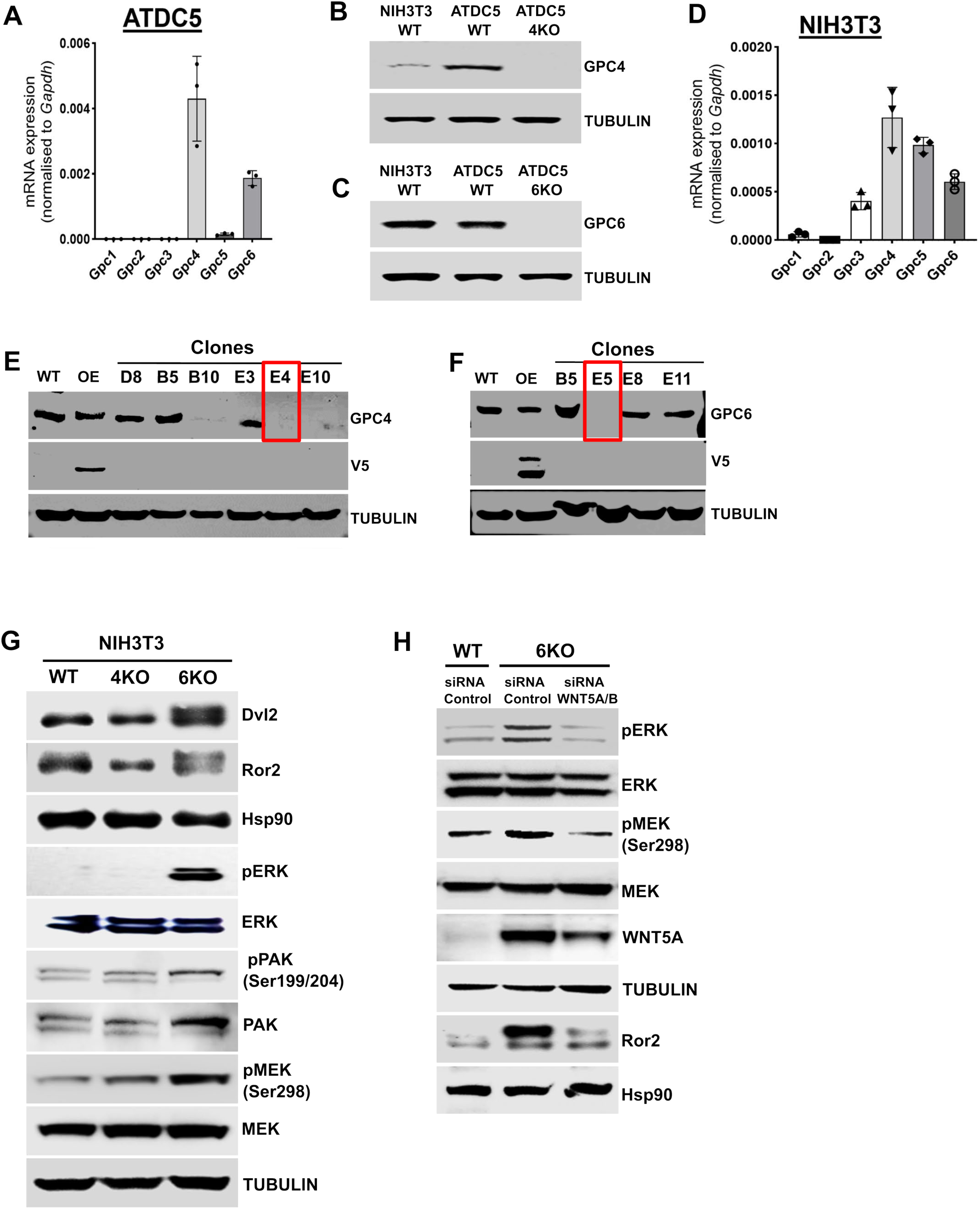
Validation of Glypican-Deficient Cell Models and Analysis of Non-Canonical WNT Signalling. **(A)** RT-qPCR analysis of glypican family member expression in ATDC5 cells, data represent mean ± SD. **(B)** Immunoblot validation of Gpc4-deficient ATDC5 cells. **(C)** Immunoblot validation of Gpc6-deficient ATDC5 cells. **(D)** RT-qPCR analysis of glypican family member expression in NIH-3T3 cells, data represent mean ± SD. **(E)** Immunoblot screening of CRISPR-edited NIH-3T3 clones identifying Gpc4-deficient cells. Red box indicates the clone selected for subsequent experiments. **(F)** Immunoblot screening of CRISPR-edited NIH-3T3 clones identifying Gpc6-deficient cells. Red box indicates the clone selected for subsequent experiments. **(G)** Immunoblot analysis of DVL2 and ROR2 together with PAK, MEK, and ERK pathway activation in WT, Gpc4-deficient, and Gpc6-deficient NIH-3T3 cells. **(H)** Immunoblot analysis of WNT5A, ROR2, and downstream signalling in WT and Gpc6-deficient NIH-3T3 cells following treatment with control or Wnt5a/b siRNA.

**Figure S4.**
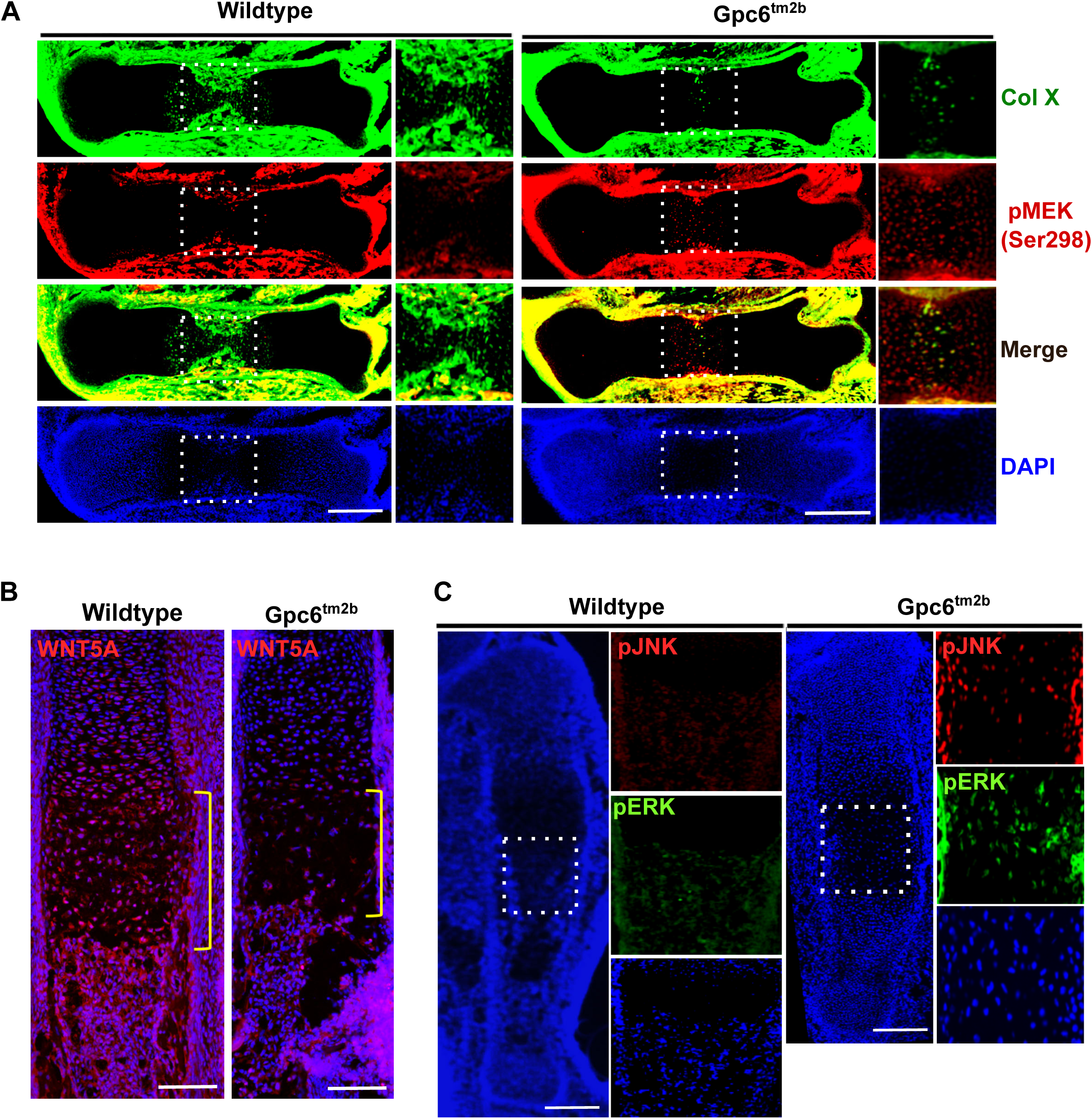
Spatial Activation of the RAC-PAK-MEK-ERK Pathway in Gpc6-Deficient Femoral and Tibial Growth Plates. **(A)** Representative immunofluorescence images of COLX and pMEK (Ser298) in E15.5 WT and *Gpc6^tm2b^* femoral growth plates. Boxed regions are shown at higher magnification and reveal increased pMEK (Ser298) signal within the prehypertrophic domain following Gpc6 loss. Scale bar 100 µm. **(B)** Representative WNT5A immunofluorescence images of E15.5 WT and *Gpc6^tm2b^* tibial growth plates, showing collapse of the physiological WNT5A gradient following Gpc6 loss. Scale bar 100 µm. **(C)** Representative immunofluorescence images of pJNK and pERK in E15.5 WT and *Gpc6^tm2b^*tibial growth plates. Boxed regions are shown at higher magnification and demonstrate coincident activation of JNK and ERK signalling within the WNT5A-producing prehypertrophic domain following Gpc6 loss. Scale bar 100 µm.

**Figure S5.**
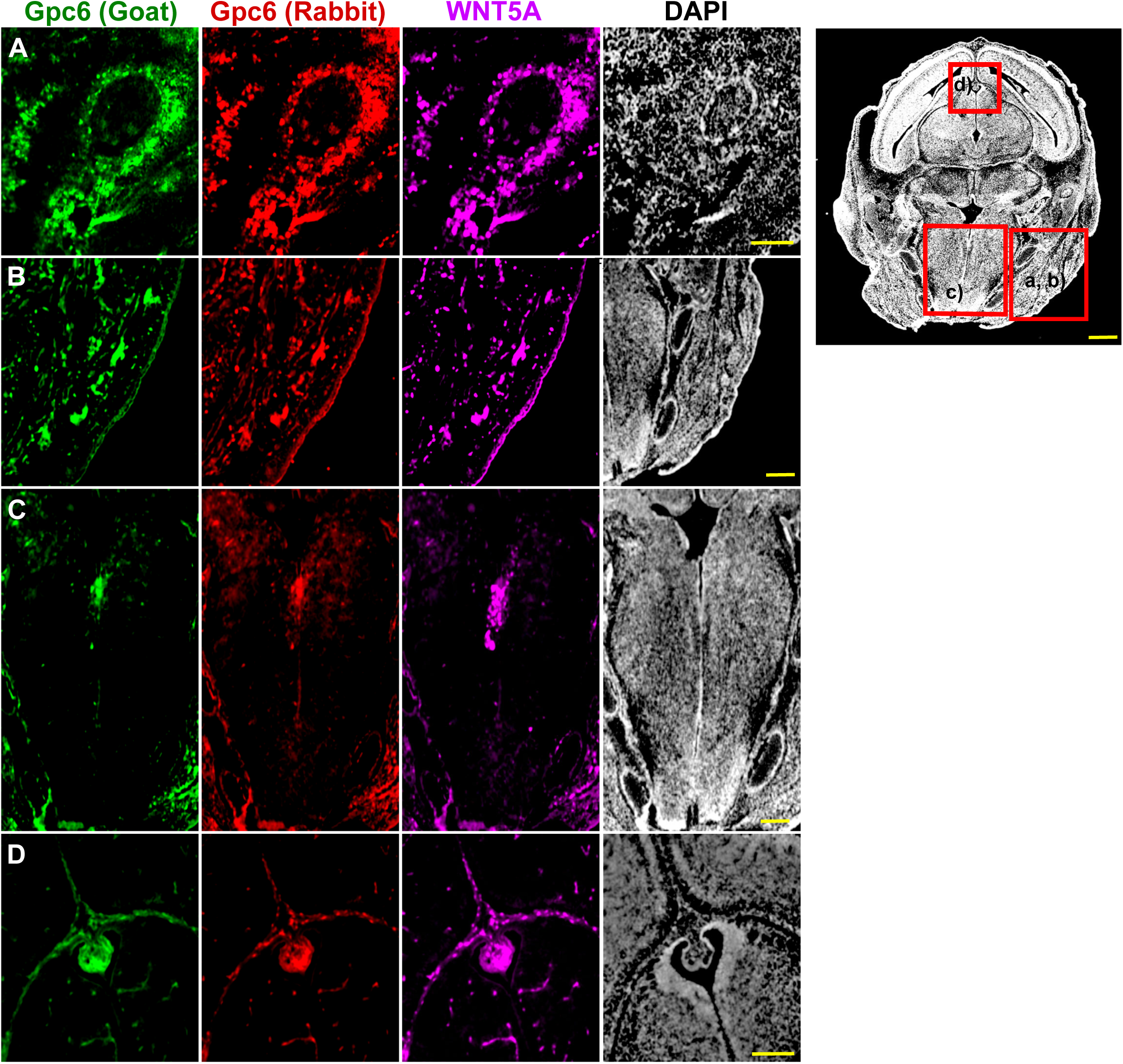
GPC6 and WNT5A Display Spatially Correspondent Expression Patterns During Embryonic Development. Representative immunofluorescence images showing GPC6 detected using independent goat- and rabbit-derived antibodies together with WNT5A staining in developing embryonic tissues. Dashed red boxes on the reference section indicate the anatomical regions analysed: **(A)** tooth bud, **(B)** whisker follicle, **(C)** tongue, and **(D)** pituitary. GPC6 and WNT5A exhibit overlapping expression patterns across each of the tissues examined. Scale bar 50 µm.

**Table S1. List of primary antibodies used in experiments**

**Table S2. Oligonucleotide sequences. (A)** Primers used for qRT-PCR analysis and **(B)** sgRNA target sequences for CRISPR/Cas9 genome editing

