## Supplemental Table 1 for "GPC6 Lipid Shielding Sustains WNT5A Gradients to Prevent a RASopathy-like Developmental State"

**Table S1. List of primary antibodies used in experiments.**

| **Antibody** | **Catalog Number** | **Source** | **Host** | **Dilution** |
| --- | --- | --- | --- | --- |
| WNT5A | MAB645 | R&D Systems | Rat | 1:200 |
| GPC6 |  |  | Rabbit | 1:500 |
| GPC6 | AF2845 | R&D Systems | Goat | 1:500 |
| Phospho-ERK1/2 (Thr202/Tyr204) | 28733-1-AP | proteintech | Rabbit | 1:500 |
| Phospho-ERK1/2 (Thr202/Tyr204) | 9101 | cell signaling | Mouse | 1:1000 |
| phospho-SAPK/JNK (Thr183/Tyr185) | 60666-1-Ig | proteintech | Mouse | 1:5000 |
| p44/42 MAPK (Erk1/2) Antibody | 9102 | cell signaling | Rabbit | 1:1000 |
| SAPK/JNK Antibody | 9252 | cell signaling | Rabbit | 1:1000 |
| phospho-MEK1/2 (Ser298) | 68047-1-Ig | proteintech | Mouse | 1:5000 |
| Collagen X | ab58632 | Abcam | Rabbit | 1:1000 |
| Dvl2 | 3216 | cell signaling | Rabbit | 1:1000 |
| Phospho-Histone H3 (Ser10) | 9701 | cell signaling | Rabbit | 1:200 |
| Phospho-MEK1/2 (Ser217/221) | 9154 | cell signaling | Rabbit | 1:1000 |
| Phospho-PAK1 (Ser144)/PAK2 (Ser141) | 85044-1-RR | proteintech | Rabbit | 1:1000 |
| PAK1/2/3 | 2604 | cell signaling | Rabbit | 1:1000 |
| HSP90 | 4877 | cell signaling | Rabbit | 1:1000 |
