## Supplemental Table 1 for "GPC6 Lipid Shielding Sustains WNT5A Gradients to Prevent a RASopathy-like Developmental State"

**Table S2. Oligonucleotide sequences. (A) Primers used for qRT-PCR analysis and (B) sgRNA target sequences for CRISPR/Cas9 genome editing**

**(A)**

|  | **Forward primer (5′-3′)** | **Reverse primer (5′-3′)** |
| --- | --- | --- |
| GAPDH | TGTGTCCGTCGTGGATCTGA | CCTGCTTCACCACCTTCTTGA |
| GPC1 | CCTCTCATCAGAGCTTCTCC | AGCATATCAGCAAGAAAACCC |
| GPC2 | CTCAGCTCTTCTCGCATTC | TTCTCATAGTAGTCCCGCAG |
| GPC3 | TCAGTTCCAGTCTCCATCAG | AGCAAAGGCGATGAAGAAG |
| GPC4 | GAAACTGAAGCTCCAGGTTAC | CACGGACACTTTGCTTACTAC |
| GPC5 | TCCTCTTCCTCTTCTTCCAC | CGCCGACTGTAAGTCTAAAG |
| GPC6 | CCTGGAAGAGATGCTCAATG | TCCTCGCTGAAGTGATACTG |
| WNT5A | CTCCTTCGCCCAGGTTGTTATAG | CATTGGAGAAGGTGCGAAGACA |
| SOX9 | TCTGGAGGCTGCTGAACGA | CGTTCTTCACCGACTTCCT |
| Col X | CATAAAGGGCCC ACTTGCTA | CAGGAATGCCTTGTTCTCCT |
| Col 2a1 | CGGTCC TACGGTGTCAGG | GCAGAGGACATTCCCAGTGT |

**(B)**

|  | **Forward primer (5′-3′)** | **Reverse primer (5′-3′)** |
| --- | --- | --- |
| g GPC6 | CACCGGCTAAAGCGGTACTACACAG | AAACCTGTGTAGTACCGCTTTAGCC |
| g GPC4 | CACCGTCTGGAGCGCATGTTTCGCC | AAACGGCGAAACATGCGCTCCAGAC |
| GPC6_Target | CGCAGAAAAGTCCCTAAATGAC | ACTGAGGGTTAATCAGCTGGAA |
| GPC4_Target | CGTGAAGACATATGGCCACTTA | AAAGGCTTCAGCTGCTCTGTAT |
| U6 | GAGGGCCTATTTCCCATGATTCC |  |
